# The genetic architecture of sporadic and recurrent miscarriage

**DOI:** 10.1101/575167

**Authors:** Triin Laisk, Ana Luiza G Soares, Teresa Ferreira, Jodie N Painter, Samantha Laber, Jonas Bacelis, Chia-Yen Chen, Maarja Lepamets, Kuang Lin, Siyang Liu, Iona Y Millwood, Avinash Ramu, Jennifer Southcombe, Marianne S Andersen, Ling Yang, Christian M Becker, Scott D Gordon, Jonas Bybjerg-Grauholm, Øyvind Helgeland, David M Hougaard, Xin Jin, Stefan Johansson, Julius Juodakis, Christiana Kartsonaki, Viktorija Kukushkina, Lifelines Cohort Study, Penelope A Lind, Andres Metspalu, Grant W Montgomery, Andrew P Morris, Preben B Mortensen, Pål R Njølstad, Dale R Nyholt, Margaret Lippincott, Stephanie Seminara, Andres Salumets, Harold Snieder, Krina Zondervan, Zhengming Chen, Donald F Conrad, Bo Jacobsson, Liming Li, Nicholas G Martin, Benjamin M Neale, Rasmus Nielsen, Robin G Walters, Ingrid Granne, Sarah E Medland, Reedik Mägi, Deborah A Lawlor, Cecilia M Lindgren

## Abstract

Miscarriage is a common complex trait that affects 10-25% of clinically confirmed pregnancies^1,2^. Here we present the first large-scale genetic association analyses with 69,118 cases from five different ancestries for sporadic miscarriage and 750 cases of European ancestry for recurrent miscarriage, and up to 359,469 female controls. We identify one genome-wide significant association on chromosome 13 (rs146350366, minor allele frequency (MAF) 1.2%, *P*_meta_=3.2× ^-8^ (CI) 1.2-1.6) for sporadic miscarriage in our European ancestry meta-analysis (50,060 cases and 174,109 controls), located near *FGF9* involved in pregnancy maintenance^3^ and progesterone production^4^. Additionally, we identified three genome-wide significant associations for recurrent miscarriage, including a signal on chromosome 9 (rs7859844, MAF=6.4%, *P*_meta_=1.3× ^-8^ in controlling extravillous trophoblast motility^5^. We further investigate the genetic architecture of miscarriage with biobank-scale Mendelian randomization, heritability and, genetic correlation analyses. Our results implicate that miscarriage etiopathogenesis is partly driven by genetic variation related to gonadotropin regulation, placental biology and progesterone production.

---

Miscarriage is defined by the World Health Organisation as the spontaneous loss of an embryo or fetus weighing less than 500 grams, up to 20-22 weeks of gestation^6^. It is the most common complication of pregnancy^1,2^ and the majority of miscarriages, both sporadic (1-2 miscarriages) or recurrent (≥3 consecutive miscarriages)^7,8^, happen in the first trimester^8,9^. Miscarriage is associated with excessive bleeding, infection, anxiety, depression^10^, infertility^11^ and an increased lifetime risk of cardiovascular disease^12,13^.

The risk of miscarriage increases with maternal age^1^, and has been associated with a range of causes; embryo and oocyte aneuploidy, parental chromosomal abnormalities, maternal thrombophilias, obesity, and endocrine and immunological dysregulation^7^ but causal underlying factors remain largely unknown. Miscarriage has a genetic component^14,15^, with most studies focusing on associations of maternal genetic variants with recurrent miscarriage. A recent systematic review illustrates the small sample sizes of these studies (vast majority <200 cases) and the heterogeneous definition of cases, and as a consequence identified largely inconsistent results^16^.

To discover and map the maternal genetic susceptibility and underlying biology of sporadic and recurrent miscarriage, we combined genome-wide association results of up to 69,118 cases from different ancestries (European, Chinese, UK South-Asian, UK African, African American, Hispanic American, UK Caribbean) for sporadic miscarriage, and subsequently the results of 750 cases of European ancestry for recurrent miscarriage in the largest genetic study of miscarriage to date (**Supplementary Table 1**). While the current guidelines define recurrent miscarriage as the loss of ≥2 pregnancies before 24 weeks^17^, we defined recurrent miscarriage as having had ≥3 self-reported miscarriages^8,18^, or the ICD-10 diagnosis code N96 for habitual abortion, in order to capture the more severe end of the phenotypic distribution and to differentiate it from sporadic miscarriage, and potentially identify any differences in the underlying genetic architecture for these two conditions.

We first performed a trans-ethnic GWAS meta-analysis for sporadic miscarriage, including genotype data for 69,118 cases and 359,469 female controls (**Supplementary Tables 1 and 2, Supplementary Data**). Association summary statistics were aggregated using trans-ethnic meta-regression implemented in the MR-MEGA software^19^ for GWAS meta-analysis (**Supplementary Data**). After post GWAS filtering for variants present in at least half (n=11) of the 21 datasets, the trans-ethnic GWAS meta-analysis of 8,664,066 variants revealed a genome-wide significant locus on chromosome 7 (lead signal rs10270417, MAF=1.7%, *P*_meta_=6.0 × 10^−9^; **Supplementary Table 3**, **Supplementary Figure 1**), driven by the Kadoorie Chinese-ancestry cohort (OR_EUR_=1.0 (0.9 − 1.0); OR_Kadoorie_=86.1 (21.1 − 350.3)), and near a previously reported endometriosis susceptibility locus^20^. Since it is known that the software used for cohort-level association testing in the China Kadoorie biobank (BOLT-LMM) can overestimate significance for rare SNPs (MAF<1%) if the case fraction is <10%^21^ (MAF_Kadoorie_=0.04%, case fraction 8.9%), and the variant was absent from other Chinese-ancestry cohorts (BGI and UKBB_CHI_) due to low MAF, the variant was not taken forward for further analysis and interpretation. A population-specific effect cannot be ruled out but would require local replication.

**Figure 1:**
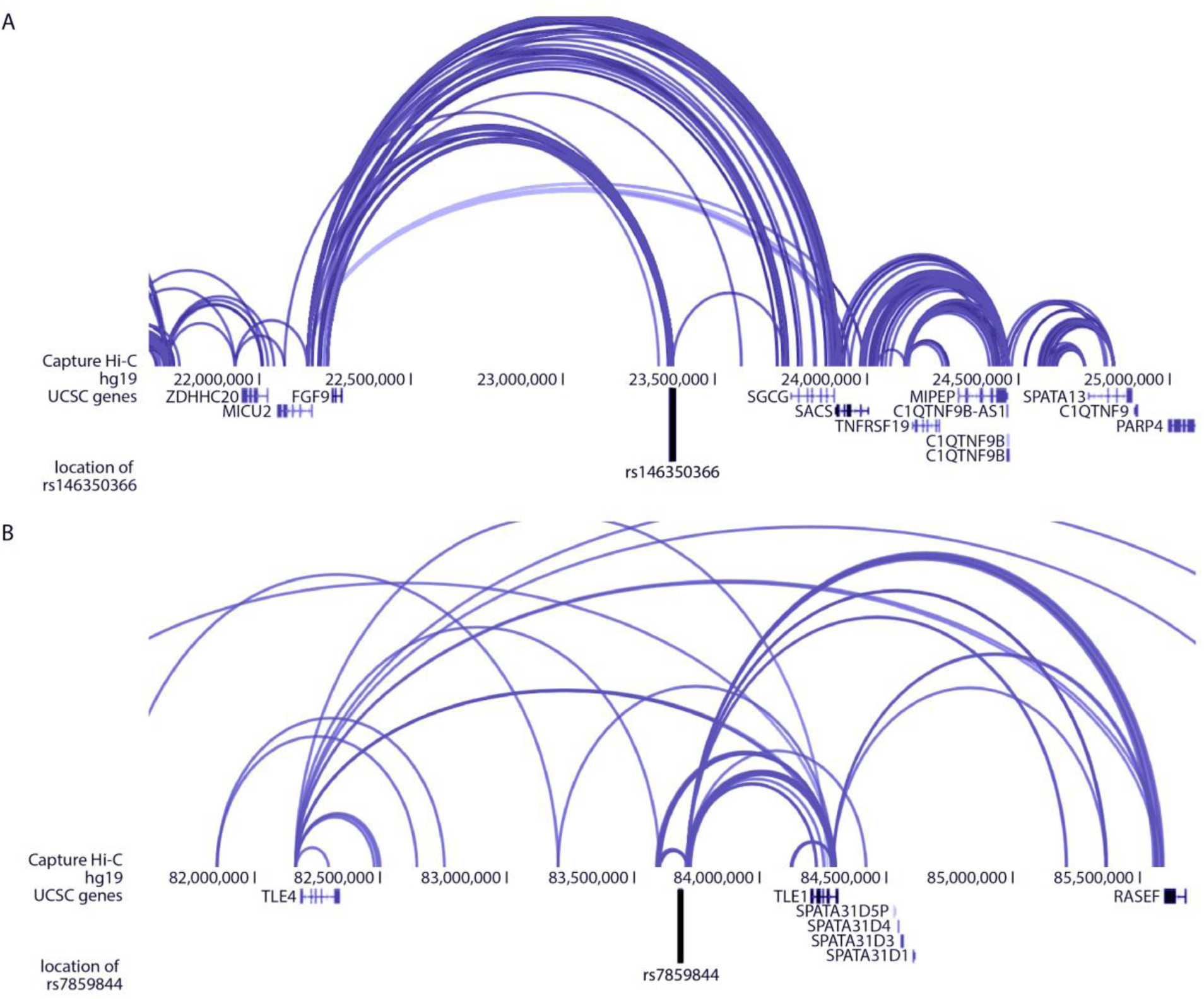
A The GWAS association rs146350366 on chromosome 13 for sporadic miscarriage in the European ancestry meta-analysis forms functional connections to the *FGF9/MICU9* region in endothelial progenitors. The black vertical line represents the location of the signal from GWAS meta-analysis. **B The GWAS association rs7859844 on chromosome 9 for recurrent miscarriage meta-analysis forms functional connections to the *TLE1* region in fetal thymus.** The black vertical line represents the location of the signal from GWAS meta-analysis. The 3D Genome Browser^35^ was used for data visualization.

We also performed a European ancestry only meta-analysis using METAL^22^, in 50,060 sporadic miscarriage cases and 174,109 female controls. After filtering for variants present in at least half (n=7) of the 13 European ancestry cohorts (n=8,275,885 SNPs), we detected one genome-wide significant locus on chromosome 13 (rs146350366, MAF=1.2%, *P*_meta_=3.2×10^−8^, OR=1.4 (1.2-1.6); **Supplementary Table 3, Supplementary Figure 2)**.

Next, we performed a European ancestry only meta-analysis aggregating summary statistics in 750 recurrent miscarriage cases and 150,215 controls from three participating cohorts (UKBB, EGCUT, ALSPAC) (**Supplementary Data**), using Stouffer’s Z-score method implemented in METAL^22^, as the effect estimates from different cohorts were not directly comparable. After filtering for variants (n=2,070,791 SNPs) with an average MAF≥0.5%, and cohort-level MAF≥0.1% as well as the same effect direction in all three cohorts, we detected four genome-wide significant signals on chromosomes 2, 9, 11, and 21 (**Supplementary Table 3**). As the initial meta-analysis approach did not yield a summary effect estimate, we applied the Firth test for significant variants to obtain uniform cohort-level association statistics and a summary effect estimate. This left us with three genome-wide significant signals: on chromosome 9 (rs7859844, MAF=6.4%, *P*_meta_=1.3×10^−8^, *P*_Firth_=2.0×10^−9^, OR=1.7 (1.4-2.0)), chromosome 11 (rs143445068, MAF=0.8%, *P*_meta_=5.2×10^−9^, *P*_Firth_=1.8×10^−10^, OR=3.4 (2.4-5.0)), and 21 (rs183453668, MAF=0.5%, Pmeta=2.8×10-8, P_Firth_=2.5×10-9, OR=3.8 (2.4-5.9)). The signal on chromosome 2 (rs138993181, MAF=0.6%, *P*_meta_=1.6×10^−8^), did not remain significant after the Firth test (*P*_Firth_=1.7×10-7, OR=3.6 (2.2-5.8)) (Supplementary Figure 3 and 4 A-D) and was not taken further for functional annotation analysis.

To our knowledge, no previous GWAS for recurrent or sporadic miscarriage have been carried out, but we checked the results for the 333 variants from a previous meta-analysis of most published idiopathic recurrent miscarriage candidate gene associations^16^ in our EUR ancestry meta-analyses for both sporadic and recurrent miscarriage. None of these variants were genome-wide significant in either the sporadic or recurrent miscarriage analysis, and only 14 (4.2%) and 11 (3%) were nominally significant (*P*<0.05) in the respective analyses (**Supplementary Table 4**). Two genome-wide linkage scans, one of 44 recurrent miscarriage cases and 44 controls and the other of 38 sibling pairs affected by idiopathic recurrent miscarriage reported loci on 6q27, 9q33.1, × p22.1^23^ and 3p12.2, 9p22.1 and 11q13.4^14^ as associated with idiopathic recurrent miscarriage but none of the recurrent or sporadic miscarriage associations identified in our much larger analysis overlap with these previously reported regions.

While previous studies have shown preliminary evidence that (recurrent) miscarriage has a genetic predisposition^14,15,23^, the heritability of miscarriage and related phenotypes has remained unquantified. We were unable to estimate the heritability for recurrent miscarriage robustly due to a relatively small number of cases. However, we estimated the traditional heritability of ‘ever having miscarried’ under a classical twin model using the QIMR twin dataset, including 1,853 monozygotic and 1,177 dizygotic complete twin pairs and 2,268 individuals from incomplete pairs, and found a heritability of 29% (95% CI 20%-38%) for any miscarriage (**Supplementary Data**, **Supplementary Table 5**). In parallel, we used the sporadic miscarriage European ancestry GWAS meta-analysis summary statistics and the LD Score regression (LDSC) method^24^ to calculate the SNP-based heritability for sporadic miscarriage. We found the SNP-based heritability estimate to be small, with h^2^=1.5% (SE 0.4) on the liability scale (assuming a population prevalence 20%). Similarly to other complex traits, our findings show the SNP-heritability is substantially lower, suggesting that other sources of genetic variation may have a larger contribution. Our study design is limited to interrogate maternal contribution to the genetic architecture of the trait, and it is likely that paternal and fetal contributions are responsible for a proportion of the heritability gap. This also prevents us from investigating the parent-offspring interaction and environmental effects, which have been shown for pre-eclampsia^25^. Overall, it might be expected that genetic factors increasing susceptibility to miscarriage are under negative selection due to their impact on reproductive fitness and hence these will be rare. Up to two-thirds of miscarriages are unrecognized and/or undiagnosed, particularly early miscarriages^26^, and thus ‘control’ women will include some misclassified as not having experienced a miscarriage. This would be expected to attenuate results towards the null and means larger numbers may be required to accurately quantify SNP-heritability and identify genome-wide significant SNPs; this is likely to affect sporadic miscarriage more so than recurrent miscarriage.

We assessed the potential genetic overlap between miscarriage phenotypes and other traits and found significant (Bonferroni corrected significance level 0.05/72=6.9×10^−4^) genetic correlations between European-ancestry sporadic miscarriage analysis and number of children (r_g_=0.69, se=0.12, *P*= 7.2×10^−9^) and age at first birth(r_g_ = −0.40, se=0.10, *P* =3.3×10^−5^)(**Supplementary Table 6**). The positive genetic correlation between sporadic miscarriage and number of children is consistent with observational associations between sporadic miscarriage and greater number of live births^27^.

We also examined associations of sporadic and recurrent miscarriage with ICD-coded disease outcomes from linked hospital episode statistics in the UKBB dataset, adjusting for number of live births and woman’s age and using FDR corrected *P*-values. We focused only on outcomes with at least one observation among the cases, resulting in testing >6,000 ICD codes for sporadic and >1,000 ICD codes for recurrent miscarriage. For sporadic miscarriage, the majority of associations were related to pregnancy, childbirth and the puerperium (*P*-values ranging between 9.9 × 10^−79^ and 4.4 × 10^−2^; **Supplementary Table 7**), supporting the observation that having more live births is associated with miscarriage^27^. Sporadic miscarriage was also positively associated with a wide variety of diagnoses, including asthma (*P*=1.6 × 10^−20^, OR=1.2 (1.19-1.3)), stillbirth (*P*=5.1 × 10^−5^,OR=74.3 (10.0-549.2)), depressive episodes (P=1.4×10^−7^, OR=1.2 (1.1-1.3)), irritable bowel syndrome (*P*=3.5×10^−9^, OR=1.3 (1.2-1.4)), intentional self-poisoning (*P*=9.5×10^−4^, OR=1.6 (1.2– 2.0)) and negatively associated with endometrial cancer (*P* =9.9×10^−3^, OR=0.8 (0.7-1.0)). Recurrent miscarriage was positively associated with tubulointerstitial nephritis (*P*=7.8×10^−5^, OR=5.3 (2.3-12.0)), infertility (P=1.9×10^−18^, OR=7.5 (4.8-11.7)), ectopic pregnancy (P=6.7×10^−17^, OR=25.4 (12.1-53.4)), and others (**Supplementary Table 8**). For some of these diagnoses, including irritable bowel syndrome, asthma, endometrial cancer, self-harm, and ectopic pregnancy, similar epidemiologic associations have been reported previously^28–32^, warranting further investigation into underlying mechanisms and highlighting the potential of large biobanks for analyzing miscarriage-associated health risks.

We also conducted a hypothesis generating phenome-wide Mendelian randomization analysis of recurrent miscarriage (using a per allele genetic risk score from the GWAS significant SNPs) in relation to 17,037 diseases and health related traits using the PHESANT^33^ package in UKBB (n=168,763) (**Supplementary Figure 6; Supplementary Data**). Three outcomes (related to alcoholism, traumatic experience, and employment history) reached Bonferroni corrected levels of statistical significance (P<2.9×10^−6^)(**Supplementary Figure 7, Supplementary Table 9**). However, these were single items from categories that include several related terms, with none of the other terms reaching suggestive thresholds of statistical significance. The failure to identify any robust causal associations between recurrent miscarriage and >17,000 variables is likely to be a combination of only having a weak genetic instrument (McFadden’s adjusted R2=0.0006, Efron’s R2=0.0002, Pseudo R2=0.0062). As the genetic instrument included only 4 rare variants we would not have been able to robustly exclude pleiotropy had effects been suggested (masking pleiotropy is possible and might contribute to null findings).

For the sporadic miscarriage European ancestry meta-analysis signal on chromosome 13, a total of five candidate SNPs and 47 potentially causal genes (13% of them protein coding) were suggested by chromatin interaction data from 21 different tissues/cell types, while no significant eQTL associations were detected using FUMA^34^ (**Supplementary Tables 10 and 11**). Capture Hi-C data from endothelial progenitor cells^35^ showed connections between the GWAS meta-analysis association signal and the *FGF9/MICU9* locus (**Figure 1A**). FGF9 plays a role in embryo implantation/pregnancy maintenance^3^, in progesterone production^4^, and has been found to be upregulated in the endometrium of women with recurrent miscarriage^36^. However, the sporadic miscarriage signal on chromosome 13 was not significant in our recurrent miscarriage meta-analysis.

For the recurrent miscarriage association signal on chromosome 9, 53 candidate SNPs and a total of 50 candidate genes were identified by chromatin interaction data: among them protein-coding genes *TLE1*, *TLE4*, *PSAT1*, *IDNK*, *GNAQ*, *RASEF*, *SPATA31D1* and *FRMD3* (**Supplementary Tables 12 and 13**). Hi-C data in fetal thymus^35^ showed interactions between the associated locus and *TLE1* (and between *TLE1* and *TLE4*) (**Figure 1B**). Both TLE1 and TLE4 are repressors of the canonical WNT signaling pathway, and participate in controlling extravillous trophoblast motility^5^. Additionally, there is evidence for the same co-repressors also regulating GnRH expression^37^. Further, a homologous protein, TLE6 has been shown to be associated with early embryonic lethality^38^ and is known to antagonize TLE1-mediated transcriptional repression^39^. On chromosome 11, both rs143445068 and rs140847838 were highlighted as potential candidate SNPs in the associated region located in the intron of *NAV2*. Chromatin interaction mapping highlighted another 17 candidate genes in the locus, including *DBX1*, *HTATIP2*, *E2F8*, *ZDHHC13*, *MRGPRX2* (**Supplementary Figure 9**). Finally, for association signal on chromosome 21, no other candidate SNPs in addition to the lead signal rs183453668 were identified, and a total of 10 candidate genes were suggested by chromatin interaction data, including *SIK1*, *U2AF1*, *CRYAA*, *HSF2BP*, *RRP1B* (**Supplementary Figure 10**). Taken together, mapping of potential candidate genes at associated loci identified several genes (*FGF9, TLE1, TLE4*) with a plausible biological role in miscarriage etiopathogenesis. However, the involvement of other transcripts at these loci cannot be ruled out and further functional studies are needed.

In conclusion, we identify four distinct susceptibility loci for sporadic and recurrent miscarriage, confirming a partly genetic basis, that does not seem to overlap, and which implicates novel biology through regulation of genes involved in gonadotropin regulation, placental biology and progesterone production.

## URLs

UK Biobank, http://www.ukbiobank.ac.uk/; Estonian Biobank, https://www.geenivaramu.ee/en; ALSPAC (http://www.bristol.ac.uk/alspac/); China Kadoorie Biobank (http://www.ckbiobank.org/); FUMA (Functional Mapping and Annotation of Genome-Wide Association Studies), http://fuma.ctglab.nl/; LD Hub, http://ldsc.broadinstitute.org/; 3D Genome Browser, http://promoter.bx.psu.edu/hi-c/;

## Acknowledgements

T.L. is supported by European Commission Horizon 2020 research and innovation programme (project WIDENLIFE, grant number 692065); Estonian Ministry of Education and Research (grants IUT34-16 and PUTJD726); Enterprise Estonia (grant EU49695). C.M.L. is supported by the Li Ka Shing Foundation, WT-SSI/John Fell funds, Oxford, NIHR Oxford Biomedical Research Centre, Oxford, Widenlife and NIH (5P50HD028138-27). D.F.C. is supported by grants from the National Institutes of Health (R01HD078641, R01MH101810 and P51OD011092). T.F. is supported by the NIHR Biomedical Research Centre, Oxford. The research in this paper has been carried out using the UK Biobank resource (under applications 17805, 11867, and 16729). We thank the study subjects for their valuable participation. We thank the twins and their families for their participation in the QIMR study. We acknowledge with appreciation all women who participated in the QIMR endometriosis study. S.E.M. and J.N.P. were supported by NHMRC applications APP1103623 and APP1084325. We are extremely grateful to all the families who took part in the ALSPAC study, the midwives for their help in recruiting them, and the whole ALSPAC team, which includes interviewers, computer and laboratory technicians, clerical workers, research scientists, volunteers, managers, receptionists and nurses. Partners HealthCare Biobank is supported by NHGRI grant U01HG008685. The authors wish to acknowledge the services of the Lifelines Cohort Study, the contributing research centers delivering data to Lifelines, and all the study participants. A full list of acknowledgements is provided in the Supplementary Data.

## Author contributions

### Data analysis

T.L., A.L.G.S, T.F., J.N.P., S.La., J.B., C-Y.C., K.L., S.Liu, A.R.; **Coordination of cohort level analysis:** H.S., Z.C., D.F.C., B.J., L.L., N.G.M., B.N., R.N., R.G.W., S.E.M., R.M., D.A.L., C.M.L.; **Provision and/or processing of (phenotype) data**: M.L., I.Y.M., J.S., M.S.A., L.Y., C.B., S.D.G., J.B-G., Ø.H., D.M.H., X.J., S.J., J.J., C.K., V.K., P.A.L., A.M., G.W.M., A.P.M., P.B.M., P.R.N., D.R.N., M.L., S.S., A.S., K.Z., I.G.; D.A.L.; **Manuscript drafting:** T.L., A.L.G.S., R.M., D.A.L., C.M.L.; All authors contributed to the final version of the manuscript.

## Additional information

All methods, additional references, supplementary figures and tables, and associated information are available in Supplementary Data.

