## Supplementary Data for "The genetic architecture of sporadic and recurrent miscarriage"

*Study cohorts*

Descriptive statistics of the cohorts included in the sporadic and recurrent miscarriage GWAS meta-analyses are presented in **Supplementary Table 1.**

*Case definitions*

Depending on the type of data available in individual cohorts, miscarriage cases were identified as follows:

<u>Sporadic miscarriage</u>: 1 or 2 self-reported miscarriages, or ICD-10 codes O02.1 and O03 on 1 or 2 separate time-points (at least 90 days between episodes).

<u>Recurrent miscarriage:</u> (i) five or more self-reported miscarriages, one live birth, no pregnancy terminations, (ii) three or more self-reported miscarriages, no live births, no pregnancy terminations, or (iii) three or more consecutive miscarriages. The latter two criteria were used to ensure the consecutive nature of the miscarriages; (iv) ICD-10 diagnosis code N96.

*Exclusion criteria*

Where data allowed, we applied the following exclusion criteria to all cohorts (the aim of these exclusions was to mainly examine associations with idiopathic miscarriage cases and thereby increase the homogeneity of the analysed phenotype):

- women with early or late menarche (<9 or >17 years), which could indicate underlying hormonal abnormalities

- women with any of diagnoses for conditions associated with increased susceptibility to miscarriage (maternal chromosomal abnormalities, thyroid conditions, neoplasms affecting endocrine glands, thrombophilias, disorders affecting the endocrine system, congenital malformations of genital organs)

*UKBB*

The UK Biobank (UKBB) is a prospective cohort of 502,637 (~5% of the >9.2 million invited) people aged 37-73 recruited in 2006-2010 from across the UK, who completed detailed questionnaires regarding socio-demographic and lifestyle characteristics and their medical history, and had a clinical assessment. Additional information about medical conditions (both existing at baseline and occurring during follow-up) has been obtained through linking with hospital admission and mortality data. Full details of the study have been reported elsewhere[1] [PMID: 25826379]. Ethics approval for the UKBB was provided by the UK National Health Service (NHS) Research Ethics Service (11/NW/0382) and all participants provided informed written consent.

Information on miscarriages was retrieved from touchscreen questionnaire fields 2774 “Have you ever had any stillbirths, spontaneous miscarriages or terminations?” and 3839 “How many spontaneous miscarriages?”, and from hospital in-patient episode data fields 41202 and 41204 (“Diagnoses – main ICD10” and “Diagnoses – secondary ICD10”, diagnosis codes O02.1 and O03, and N96). In the UKBB, most of the data has been collected during the initial assessment visit, however, for some participants, data has been additionally collected on repeat assessment visits. Therefore, if a participant has answered the same question on multiple occasions, answers were aggregated, excluding participants who have given discordant answers on different occasions. Participants giving no answer to relevant questions, or answering “Prefer not to answer” or “Do not remember” were excluded from the analysis. Analyses were performed under data applications 17805 (“Dissemination of shared genetics across phenotypes associated with reproductive health and related endophenotypes”), 11867 (“Dissection of the Genetic Susceptibility of Obesity Traits and their Comorbidities”), and 16729 (“MR-PheWAS: hypothesis prioritization among potential causal effects of body mass index on many outcomes, using Mendelian randomization”). The GWAS analysis included 37,150 women of White European ancestry with self-reported or electronic health record-derived sporadic miscarriage, 421 with recurrent miscarriage and 164,775 female controls (no miscarriages and no exclusion diagnoses) with available genome-wide data. The sporadic miscarriage trans-ethnic meta-analysis also included 511 cases and 1,424 controls of UK South-Asian ancestry, 390 cases and 957 controls of UK Caribbean ancestry, 132 cases and 433 controls of UK Chinese ancestry, and 273 cases and 482 controls of UK African ancestry.

*EGCUT*

The Estonian Genome Center of the University of Tartu (EGCUT; http://www.biobank.ee) is a population-based biobank with a total cohort size of 51,515 participants (aged 18-85+)[2]. The EGCUT cohort included a total of 3,368 women of White European ancestry with sporadic miscarriage, 113 with recurrent miscarriage, and 17,996 women as controls. Information on miscarriages was retrieved from questionnaire fields “How many times have you got pregnant?” and “How many of the pregnancies ended with unintentional miscarriage?”, and based on ICD codes obtained from linking with Health Insurance Fund databases. The study was approved by the Ethics Review Committee of the University of Tartu (243T-12). All biobank participants have additionally signed a broad informed consent form.

*ALSPAC*

The Avon Longitudinal Study of Parents and Children (ALSPAC) is a prospective pregnancy/birth cohort that recruited 14,541 pregnancies of women resident in Avon, UK with expected dates of delivery 1st April 1991 to 31st December 1992. These women delivered 14,062 live births and they, their partners and offspring have been followed-up since then with detailed repeat questionnaires, hands-on clinic assessments and record linkage. Full details of the study can be found elsewhere[3]. ALSPAC is an accessible resource for the research community and the study website contains details of all the data that are available through a fully searchable data dictionary (http://www.bris.ac.uk/alspac/researchers/data-access/data-dictionary). Ethical approval for the ALSPAC was obtained from the ALSPAC Ethics and Law Committee and the UK National Health Service Research Ethics Committee (full details at http://www.bristol.ac.uk/alspac/researchers/data-access/ethics/lrec-approvals/#d.en.164120). Women provided written informed consent.

At recruitment and follow-up the women have been asked about miscarriage and stillbirth. For this study, we considered miscarriages reported at baseline regarding pregnancies before the index child, and at subsequent follow-ups regarding pregnancies after the index child. At baseline, women were asked “Have you ever had any miscarriages?” and “How many times have you miscarried?”. At the follow-up assessments, miscarriage was retrieved from information asked about: i. outcome of each pregnancy after the index child (i.e. “Since child’s <age at follow-up>’s birthday, have you become pregnant?”, “What happened in the 1st pregnancy?”, “What happened in the 2nd pregnancy?”), ii. occurrence of miscarriage (i.e. “Since the study child was <age at follow-up>, have you had a miscarriage?”), and iii. whether a dilation and curettage (D&C) occurred due to miscarriage (i.e. “Have you had a D and C (scrape) in the last 2 years?” “Was this because of miscarriage?”). In total, 1,473 women had sporadic miscarriage, 216 had recurrent miscarriage, and 4,475 women had no miscarriage.

*QIMR*

The QIMR samples were drawn from two cohorts of adult twins and their relatives (parents, siblings, adult children and spouses) who have taken part in a wide range of studies of health and well-being via previous postal questionnaires and telephone interview studies (questionnaires and methods are summarized in Medland et al., 2008[4]). As a result, this sample includes related individuals, 1,145 women reporting sporadic miscarriage and 5,136 women who did not report experiencing miscarriage as controls. The QIMR Endo samples were drawn from a cohort of women with a confirmed surgical diagnosis of endometriosis, and for whom detailed reproductive history data are available[5,6]. This sample includes only unrelated individuals, 497 women reporting sporadic miscarriage and 1,078 women who did not report miscarriage as controls. For both samples miscarriage information was drawn from questionnaire fields “Have you ever had a miscarriage?” and “Number of miscarriages?”. Ethics approval for studies involving these individuals was granted by the QIMR Human Research Ethics Committee. Informed consent was obtained from all participants.

*iPSYCH*

The iPSYCH2012 (Lundbeck Foundation Initiative for Integrative Psychiatric Research) dataset represents a Danish case-control cohort (a total of 76,657 participants) for psychiatric disorders. The iPSYCH cohort included a total of 1,173 women with sporadic miscarriage and 4,821 women as controls (all of White European ancestry). Women were considered sporadic miscarriage cases if they had an inclusion code (O02.1 and/or O03) on one or two separate occasions in their medical record. At the same time, individuals with any of the abovementioned exclusion diagnoses were not included in the analysis. Controls were selected from women who had not had a miscarriage and were matched based on genotyping wave, four controls for each case.

*Lifelines*

The Lifelines dataset included in this study represents a subset of samples with available genotype data from the Lifelines prospective population-based cohort[7,8], examining in a unique three-generation design the health and health-related behaviours of 167,729 persons living in the North of The Netherlands. It employs a broad range of investigative procedures in assessing the biomedical, socio-demographic, behavioural, physical and psychological factors which contribute to the health and disease of the general population, with a special focus on multi-morbidity and complex genetics. The Lifelines dataset was accessed under data application OV17-0393. Self-reported information on miscarriages was extracted from questionnaire field “How many miscarriages (up to 16 week) have you had?”. The sample includes 1,676 sporadic miscarriage cases and 5,091 female controls. The Lifelines study has been approved by the review board of the University Medical Center, Groningen, and adheres to the principles expressed in the Declaration of Helsinki. All study participants provided written informed consent.

*Partners HealthCare Biobank*

The Partners HealthCare Biobank dataset represents a hospital-based biobank at Partners HealthCare (the parent organization of Massachusetts General Hospital and Brigham and Women’s Hospital) representing the U.S. general population. The Partners HealthCare Biobank maintains blood and DNA samples from consented patients seen at Partners HealthCare hospitals in the Boston area of Massachusetts. For the analyses described in this paper, only European American patients were included due to sample size. Patients are recruited in the context of clinical care appointments, and also electronically at Partners HealthCare. All patients participating in the Partners Biobank have given a consent for linking their samples to clinical information. Sporadic miscarriage cases were identified using ICD codes and the same inclusion and exclusion criteria described above, resulting in 58 cases and 289 controls for analysis.

*MoBa-HARVEST*

The Norwegian Mother and Child Cohort Study (MoBa) is a prospective population-based pregnancy cohort study conducted by the Norwegian Institute of Public Health. Participants were recruited from all over Norway from 1999-2008. The women consented to participation in 41% of the pregnancies. The cohort now includes 114 500 children, 95 200 mothers and 75 200 fathers. Blood samples were obtained from both parents during pregnancy and from mothers and children (umbilical cord) at birth. We are grateful to all the participating families in Norway who take part in this on-going cohort study. The cohort is described in the following publications[9–11]. In the current study, a subset (n=8,000) of the MoBa cohort (genotyping effort HARVEST, also known as Njolstad1) is included. Sporadic miscarriage cases were defined based on questionnaire-derived data, and controls were also restricted to have at least one previous delivery. The analysis included 1,653 sporadic miscarriage cases and 3,199 female controls.

*BGI Chinese millionome database Phase I 140K study*

The BGI cohort represents a dataset of 141,431 Chinese women generated for non-invasive prenatal testing (NIPT), for whom whole-genome low-pass sequencing data is available[12]. A subset of 135,115 participants who reported their pregnancy history were included in this study, resulting in 8,865 sporadic miscarriage cases and 126,290 controls for analysis.

*China Kadoorie Biobank*

The China Kadoorie Biobank is a prospective population-based cohort of 512,891 adults aged 30-79 years recruited from 10 geographically defined regions during 2004-2008, with collection of questionnaire data, physical measurements and blood samples[13]. The current study included data for 57,622 women, from whom 5,038 sporadic miscarriage cases and 51,696 controls who had ever been pregnant were extracted based on self-reported data on miscarriages. Local, national and international ethics approval was obtained and all participants provided written informed consent.

*Women's Health Initiative*

The Women's Health Initiative (WHI), sponsored by the National Heart, Lung, and Blood Institute (NHLBI), is a long-term national health study that focuses on strategies for preventing heart disease, breast and colorectal cancer, and osteoporosis in postmenopausal women. Our meta-analysis included data from several GWAS substudies from the WHI project: WHI HIPFX (mostly European ancestry case-control study for hip fracture; 273 sporadic miscarriage cases and 708 controls), WHI Gecco cyto (European ancestry case-control study by the Genetics and Epidemiology of Colorectal Cancer Consortium; 274 sporadic miscarriage cases and 609 controls), WHI WHIMS (WHI Memory Study; 411 sporadic miscarriage European ancestry cases and 950 controls), WHI GARNET (Genomics and Randomized Trials Network cohort study; 954 sporadic miscarriage cases and 2,096 controls of European American ancestry), WHI SHARE (SNP Health Association Resource; 1,151 sporadic miscarriage cases and 2,195 controls of Hispanic American ancestry, and 2,919 sporadic miscarriage cases and 4,546 controls of African American ancestry). Cases were identified based on self-administered questionnaire data from variable phv00078626.v6.p3 ('How many miscarriages or unspecified type of pregnancies').

**GWAS genotyping and imputation**

Details on cohort-level genotyping, quality control (QC), and imputation can be found in **Supplementary Table 2.**

**Association analyses and meta-analysis**

Details on how association analyses were carried out on the cohort level can be found in **Supplementary Table 2**. Cohort-level association analyses had been performed using genotype data imputed to suitable reference panels and adjusted for year of birth. Where available and appropriate, additional cohort-specific covariates, such as principal components or genotyping array, were used to correct for potential within-cohort stratification.

*Meta-analysis*

Central QC was conducted using EasyQC[14]. During central QC, allele frequencies and alignment were compared against suitable reference datasets (Haplotype Reference Consortium[15], 1000 Genomes[16]) to detect potential strand issues or large allele frequency deviations from the reference population. Monomorphic markers, and also markers with strand mismatch, poor imputation quality (INFO score <0.4) or an arbitrary minor allele count cut-off <=6 were excluded from each study prior to the meta-analysis. The results from individual cohorts were meta-analysed in parallel by two different analysts. All genome-wide significant variants that passed the applied filters (see below) are listed in **Supplementary Table 3**.

For the trans-ethnic meta-analysis, we used the MR-MEGA software[17], adjusting for the first two principal components. After the meta-analysis, we applied an additional filter for variants present in at least half (n=11) of the cohorts, to rule out spurious associations. This resulted in 8,664,066 variants, and a genome-wide significant association on chromosome 7 (rs10270417). Indels were not considered due to their lower quality. A closer inspection of the effect sizes for the observed association in individual cohorts revealed the association was mainly driven by one of the Chinese-ancestry cohorts (**Supplementary Figure 1E**) where the MAF was 0.04%. It is known that BOLT-LMM, used for analysis in the Kadoorie cohort, can overestimate significance for rare SNPs (MAF <1%) if the case fraction is <10%[18]; therefore we performed additional analyses. The Kadoorie samples have been collected from 10 different region centers, therefore we checked for batch effects. Adding batch ID as a covariate did not have a significant impact on the association statistics (original *P*-value $5.0\times10^{-10}$, after adding batch ID as covariate $P=2.2\times10^{-15}$). To check possible confounding effect from samples being collected from 10 different region centres, we performed separate analyses for each region centre, followed by meta-analysis. As the SNP is very rare, SNPTEST failed to converge in five research centre datasets; however, fixed effect meta-analysis detected a similar effect direction in the remaining datasets, although with significant differences in effect magnitude across different region centres ($P_{meta}$=4.8×10$^{-4}$; $P_{het}$=5.2×10$^{-5}$) and a considerably larger *P*-value compared to the BOLT-LMM results. Although the two methods (SNPTEST and BOLT-LMM) are not directly comparable, given the absence of this variant in other Chinese ancestry cohorts (BGI and $UKBB_{CHI}$), the rs10270417 signal was not taken further for functional annotation.

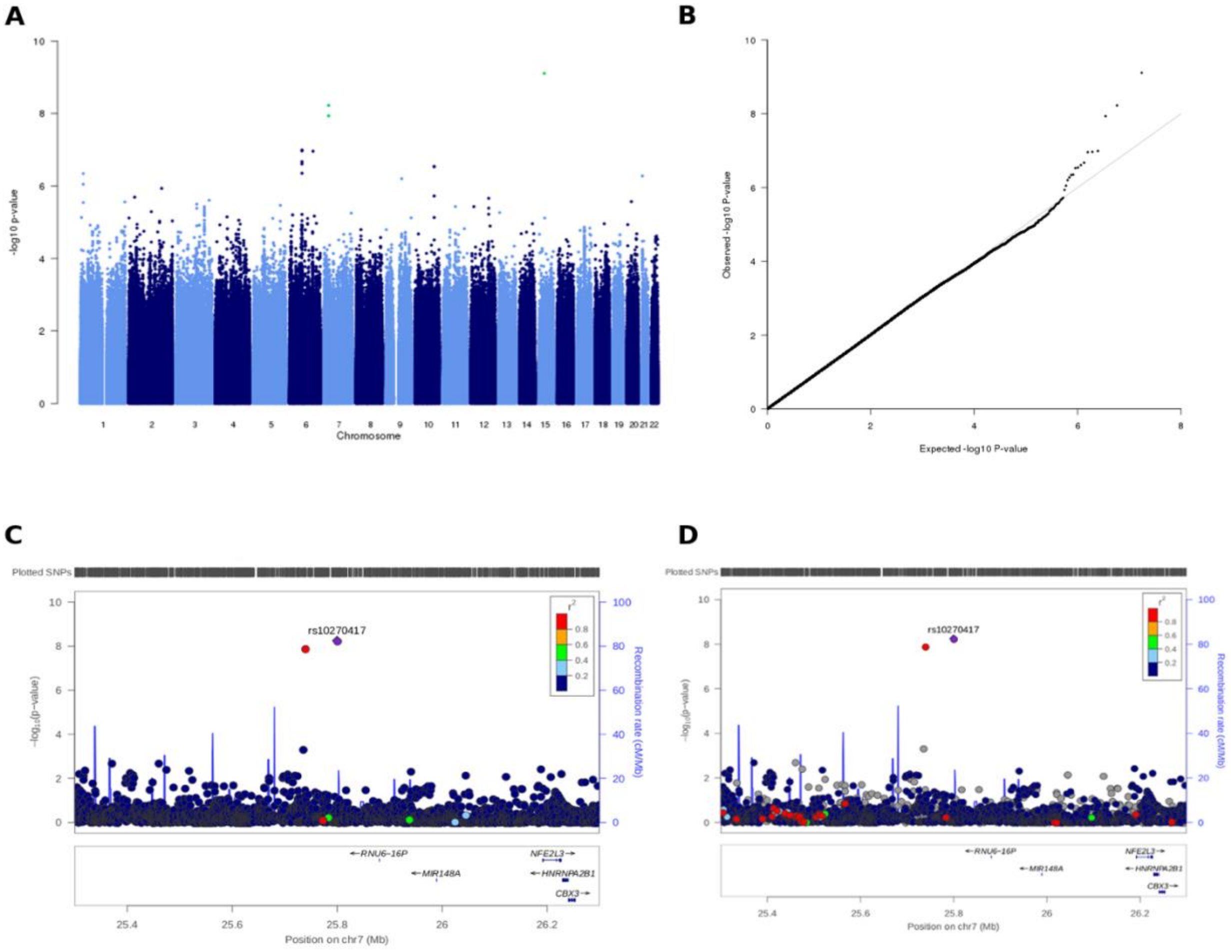

**E**

### Sporadic miscarriage trans-ethnic meta-analysis
## rs10270417

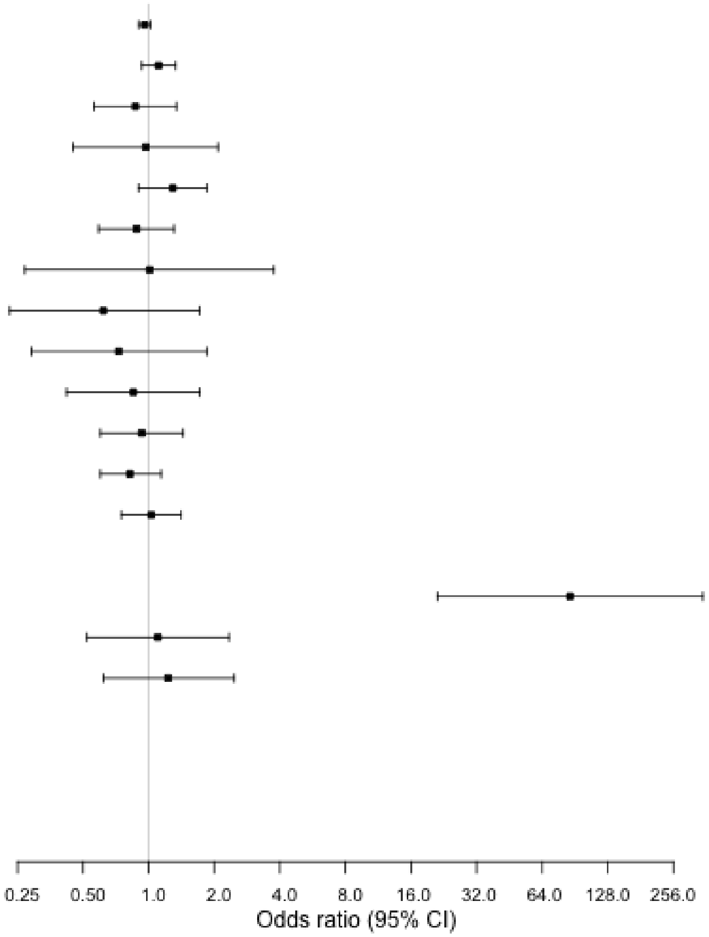


| Cohort | P-value | OR (95% CI) | N_case/N_control |
|---|---|---|---|
| UKBB | 0.22 | 0.96 (0.9-1.02) | 37,105/127,670 |
| EGCUT | 0.2512 | 1.11 (0.93-1.33) | 3,368/17,996 |
| QIMR | 0.5163 | 0.87 (0.56-1.34) | 1,145/5,136 |
| QIMREndo | 0.9402 | 0.97 (0.45-2.08) | 497/1,078 |
| iPSYCH | 0.1708 | 1.29 (0.9-1.85) | 1,173/4,812 |
| LifeLines | 0.5257 | 0.88 (0.59-1.31) | 1,676/5,091 |
| PBK | 0.9895 | 1.01 (0.27-3.76) | 58/289 |
| WHI_HIPFX | 0.341 | 0.62 (0.23-1.71) | 273/708 |
| WHI_GECCO | 0.4954 | 0.73 (0.29-1.85) | 274/609 |
| WHI_WHIMS | 0.6388 | 0.85 (0.42-1.72) | 411/950 |
| WHI_GARNET | 0.7549 | 0.93 (0.6-1.44) | 954/2,096 |
| MOBA_HARVEST | 0.2409 | 0.82 (0.6-1.14) | 1,653/3,199 |
| ALSPAC | 0.8494 | 1.03 (0.75-1.41) | 1,473/4,475 |
| BGI | | NA | 8,865/126,290 |
| Kadoorie | 5e-10 | 86.06 (21.14-350.29) | 4,817/49,033 |
| WHI_SHARE_AA | 0.8007 | 1.1 (0.52-2.34) | 2,919/4,546 |
| WHI_SHARE_HA | 0.5538 | 1.23 (0.62-2.47) | 1,151/2,195 |
| UKBB_SAS | | NA | 511/1,424 |
| UKBB_CHI | | NA | 132/433 |
| UKBB_CAR | | NA | 390/957 |
| UKBB_AFR | | NA | 273/482 |

**Supplementary Figure 1. Manhattan (A), QQ (B) and regional plots (C,D) for sporadic miscarriage trans-ethnic meta-analysis for markers present in at least half (n=11) of the cohorts.** Regional plot depicts SNPs plotted by their position and GWAS meta-analysis –log10(P-value) for association with sporadic miscarriage using the 1000G EUR (C) and ASN (D) as a reference for plotting, respectively**. (E) Forest plot of association statistics in individual cohorts**.

European-ancestry only sporadic miscarriage meta-analysis was carried out with METAL[19] using inverse variance fixed effects meta-analysis and single genomic correction. Recurrent miscarriage meta-analysis was conducted using METAL[19] and Stouffer's (*P*-value based effective sample size weighted) method and single genomic correction. After the analysis, sporadic miscarriage meta-analysis results were additionally filtered to exclude markers not present in at least half of the cohorts (n=7), while from the recurrent miscarriage meta-analysis results we excluded variants that did not have the same effect direction in all three cohorts, had an average MAF of <0.5%, and a MAF of <0.1% in any of the three cohorts. Indels were not considered due to their lower quality. The quantile-quantile plots, Manhattan plots and locus zoom plots of the meta-analyses are shown on **Supplementary Figures 2 and 3**. In the sporadic miscarriage European ancestry meta-analysis, we detected an association signal on chromosome 13 (**Supplementary Table 3**), which was taken further for functional annotation.

In the recurrent miscarriage meta-analysis, we detected four genome-wide significant signals on chromosomes 2, 9, 11, and 21 after applying the aforementioned filters (**Supplementary Table 3**). In order to obtain uniform effect estimates for the sentinel markers in these loci, the Firth test was used to recalculate cohort-level association statistics. The association on chromosome 2 did not remain significant after recalculations with the Firth test.

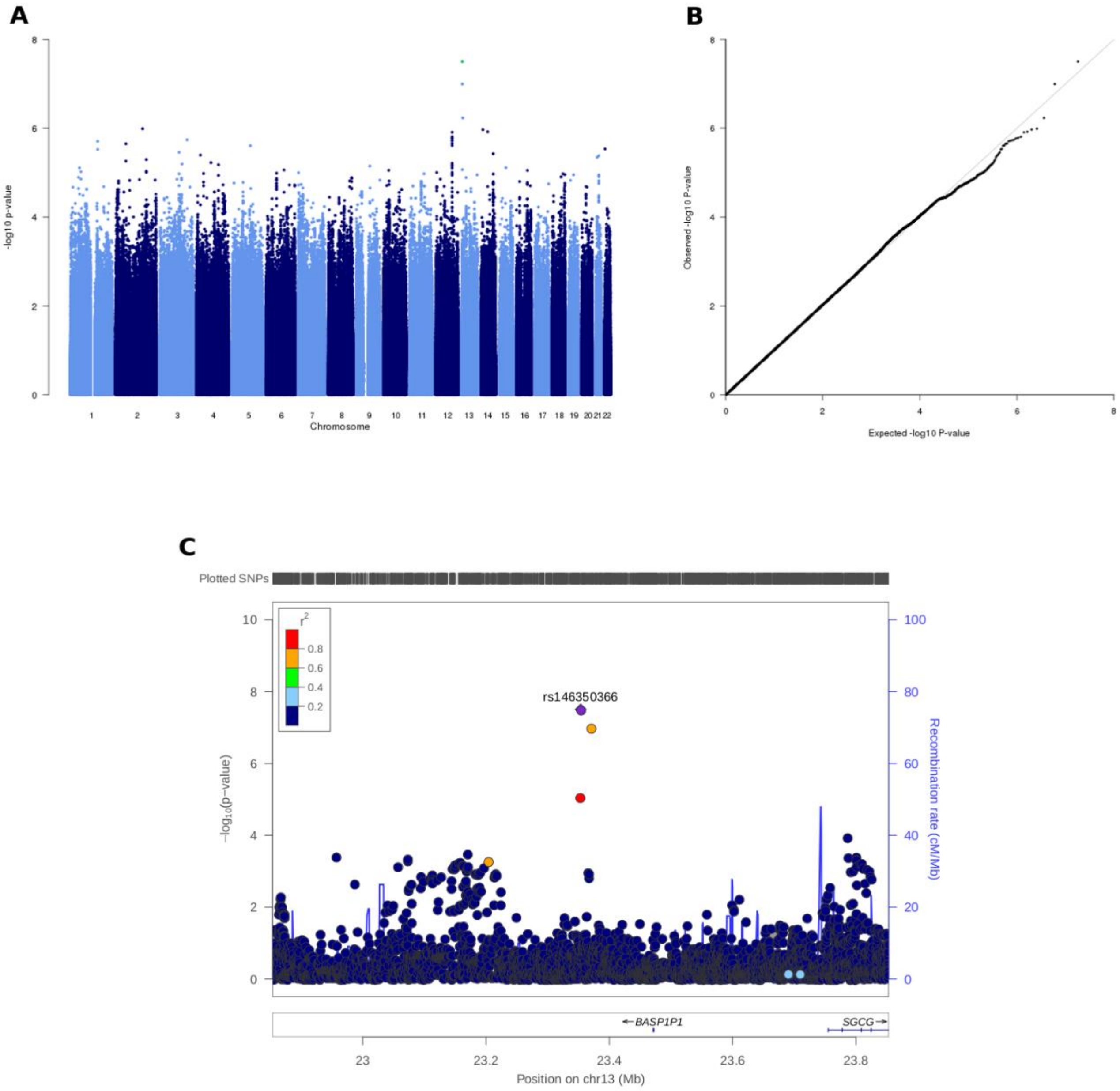
A
-log10 p-value
Chromosome
B
Observed -log10 P-value
Expected -log10 P-value
C
Plotted SNPs
r2
0.8
0.6
0.4
0.2
rs146350366
−log10(p-value)
Recombination rate (cM/Mb)
←BASP1P1
SGCG→
Position on chr13 (Mb)

**D**

**Sporadic miscarriage European ancestry meta-analysis**
**rs146350366**

| Cohort | P-value | OR (95% CI) | N_case/N_control |
|---|---|---|---|
| UKBB | 4.7e-05 | 1.38 (1.18-1.6) | 37,105/127,670 |
| EGCUT | 3e-05 | 1.51 (1.25-1.84) | 3,368/17,996 |
| QIMR | 0.96 | 1.02 (0.86-1.21) | 1,145/5,136 |
| QIMREndo | 0.87 | 1.18 (0.78-1.78) | 433/1,078 |
| iPSYCH | | NA | 1,173/4,812 |
| LifeLines | | NA | 1,676/5,091 |
| PBK | | NA | 58/289 |
| WHI_HIPFX | 0.39 | 0.51 (0.11-2.26) | 273/708 |
| WHI_GECCO | 0.57 | 0.51 (0.05-5.08) | 274/609 |
| WHI_WHIMS | 0.42 | 1.89 (0.37-9.72) | 411/950 |
| WHI_GARNET | 0.86 | 1.09 (0.42-2.84) | 954/2,096 |
| MOBA_HARVEST | 0.63 | 1.21 (0.55-2.63) | 1,653/3,199 |
| ALSPAC | 0.85 | 0.93 (0.46-1.9) | 1,473/4,475 |
| **Meta-analysis** | **3.15e-08** | **1.39 (1.24-1.56)** | **49,996/174,109** |

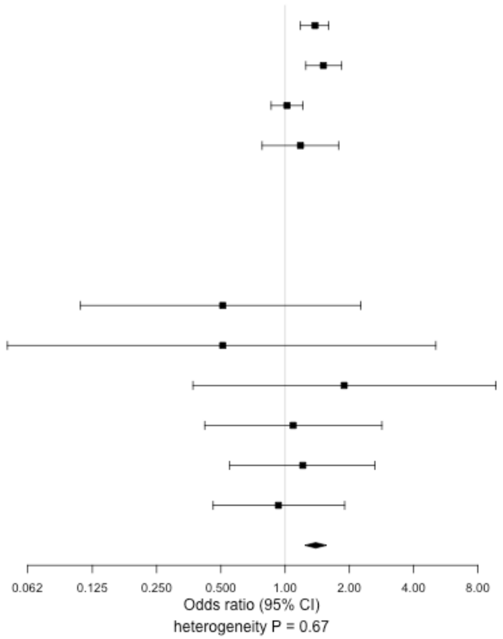


**Supplementary Figure 2. Manhattan (A), QQ (B), regional (C) and forest (D) plots for sporadic miscarriage European ancestry meta-analysis filtered for markers present in at least half (n=7) of the cohorts.** The genome-wide significant marker on chromosome 13 (rs146350366) is highlighted in green on the Manhattan plot and a close-up of the locus is shown on the regional plot in the lower panel.

A B

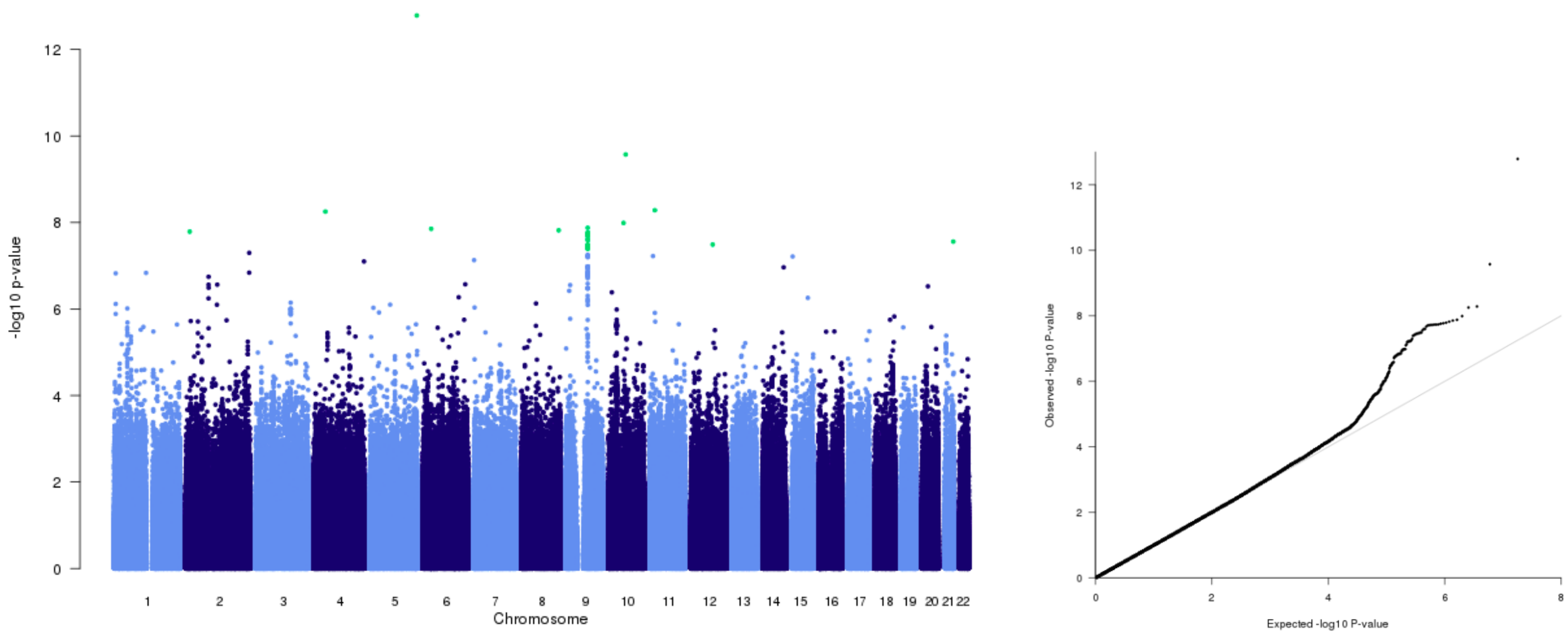


**Supplementary Figure 3. Manhattan (A) and QQ (B) plots for recurrent miscarriage European ancestry meta-analysis filtered for markers present in at least two cohorts, with and average MAF of 0.5% and cohort-level MAF of 0.1%.** Genome-wide significant loci are highlighted in green on the Manhattan plot. Subsequent filter were applied to significant loci to remove indels and those markers not present in all three cohorts.

**A**

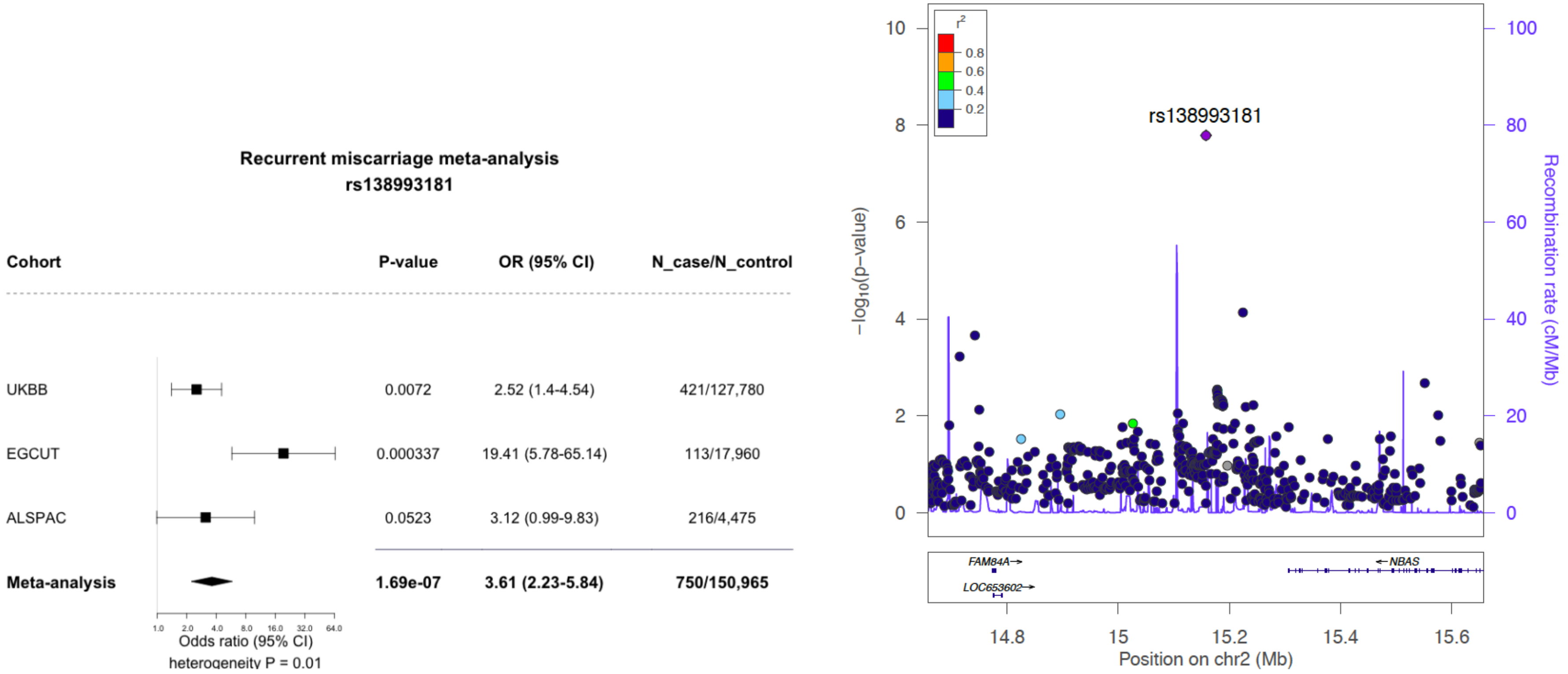
Recurrent miscarriage meta-analysis
rs138993181
Cohort
P-value
OR (95% CI)
N_case/N_control
UKBB
0.0072
2.52 (1.4-4.54)
421/127,780
EGCUT
0.000337
19.41 (5.78-65.14)
113/17,960
ALSPAC
0.0523
3.12 (0.99-9.83)
216/4,475
Meta-analysis
1.69e-07
3.61 (2.23-5.84)
750/150,965
Odds ratio (95% CI)
heterogeneity P = 0.01
rs138993181
r2
0.8
0.6
0.4
0.2
−log10(p−value)
Recombination rate (cM/Mb)
FAM84A
LOC653602
NBAS
Position on chr2 (Mb)

**B**

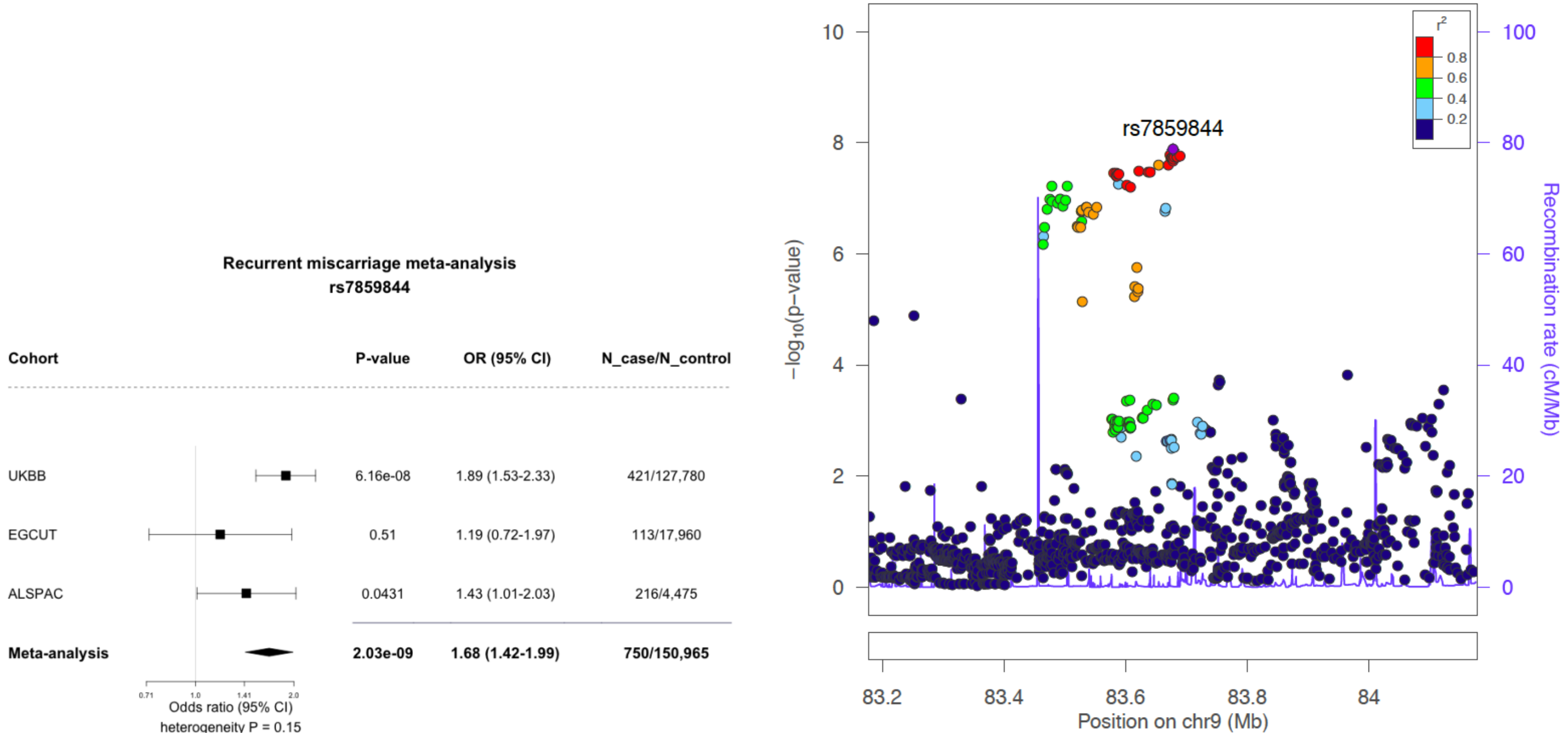
Recurrent miscarriage meta-analysis
rs7859844
Cohort
P-value
OR (95% CI)
N_case/N_control
UKBB
6.16e-08
1.89 (1.53-2.33)
421/127,780
EGCUT
0.51
1.19 (0.72-1.97)
113/17,960
ALSPAC
0.0431
1.43 (1.01-2.03)
216/4,475
Meta-analysis
2.03e-09
1.68 (1.42-1.99)
750/150,965
0.71
1.0
1.41
2.0
Odds ratio (95% CI)
heterogeneity P = 0.15
rs7859844
r²
0.8
0.6
0.4
0.2
−log10(p−value)
Recombination rate (cM/Mb)
10
8
6
4
2
0
100
80
60
40
20
0
83.2
83.4
83.6
83.8
84
Position on chr9 (Mb)

C

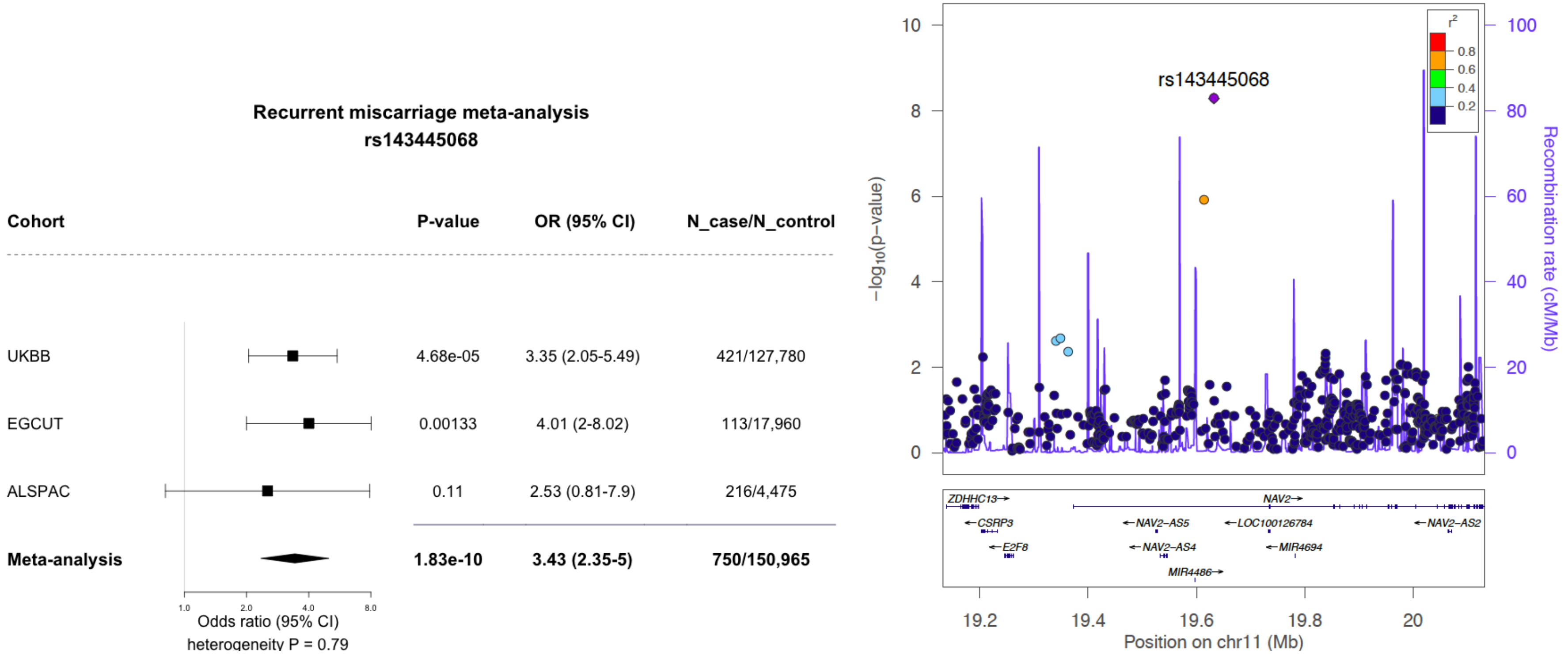
Recurrent miscarriage meta-analysis
rs143445068
Cohort
P-value
OR (95% CI)
N_case/N_control
UKBB 4.68e-05 3.35 (2.05-5.49) 421/127,780
EGCUT 0.00133 4.01 (2-8.02) 113/17,960
ALSPAC 0.11 2.53 (0.81-7.9) 216/4,475
Meta-analysis 1.83e-10 3.43 (2.35-5) 750/150,965
1.0
2.0
4.0
8.0
Odds ratio (95% CI)
heterogeneity P = 0.79
rs143445068
−log10(p−value)
Recombination rate (cM/Mb)
r2
0.8
0.6
0.4
0.2
ZDHHC13
CSRP3
E2F8
NAV2
NAV2−AS5
NAV2−AS4
MIR4486
LOC100126784
MIR4694
NAV2−AS2
19.2
19.4
19.6
19.8
20
Position on chr11 (Mb)

**D**

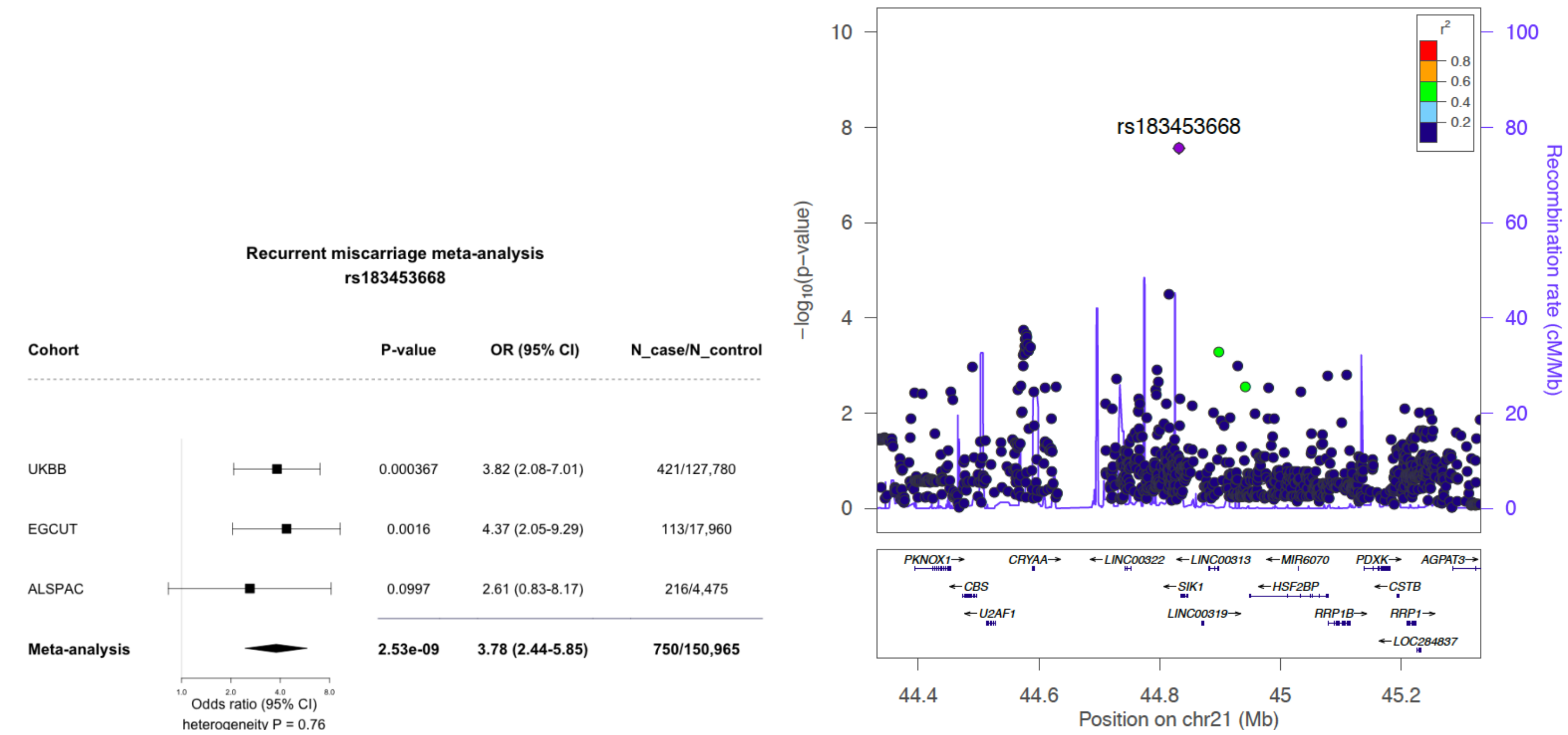


**Supplementary Figure 4. Regional and forest plots for recurrent miscarriage European ancestry meta-analysis top hits.** Effect size and summary estimates from Firth test. A) rs138993181 on chromosome 2; B) rs7859844 on chromosome 9; C) rs143445068 on chromosome 11; D) rs183453668 on chromosome 21.

### Gene-based analyses

Gene-based genome-wide association analysis was carried out with MAGMA 1.6[20] (Multi-marker Analysis of GenoMic Annotation) with default settings implemented in FUMA[21]. Briefly, variants located in the gene body were assigned to respective protein-coding genes (n=18,929;

Ensembl build 85), and the resulting SNP *P*-values are combined into a gene test-statistic using the SNP-wise mean model[20]. To adjust for multiple testing, genome-wide significance level was set at $2.6\times10^{-6}$, according to the number of tested genes. No genes passed the threshold of significance.

**Look-up of variants previously associated with recurrent miscarriage**

We conducted a lookup in our summary statistics of all variants included in a recent, extensive and systematic review / meta-analysis of published genetic association studies in idiopathic recurrent spontaneous abortion[22]. The results of this lookup are given in **Supplementary Table 4**.

**Heritability analysis**

The sporadic miscarriage GWAS European-ancestry meta-analysis summary statistics and LD Score Regression (LDSC) method[23] were used for heritability estimation. The linkage disequilibrium (LD) estimates from European ancestry samples in the 1000 Genomes projects were used as a reference. Heritability estimates were converted to the liability scale using a population prevalence of 0.2 for sporadic miscarriage. Using the UKBB SNP-Heritability Browser (https://nealelab.github.io/UKBB_ldsc/h2_browser.html), we also did a look-up for different versions of the miscarriage phenotype or related phenotypes in the UKBB dataset and observed similar heritability estimates for 'Ever had stillbirth, spontaneous miscarriage or termination'($h^2$=0.04; s.e.=0.008; population prevalence 31.5%) and 'Number of spontaneous miscarriages' ($h^2$=0.03; s.e.=0.01).

Data from 1,853 complete female monozygotic (MZ) and 1,177 dizygotic (DZ) twin pairs and 2,268 women from incomplete or opposite sex twin pairs (mean year of birth 1954, range 1893-1989) from the QIMR dataset were used to estimate heritability under a classical twin model, using a multifactorial threshold model in which discrete traits are assumed to reflect an underlying normal distribution of liability (or predisposition). Liability, which represents the sum of all the multifactorial effects, is assumed to reflect the combined effects of a large number of genes and environmental factors each of small effect[24]. All data analyses were conducted using maximum likelihood analyses of raw data within Mx[25]. Corrections for year of birth were included with the model, such that the trait value for individual *j* from family *i* was parameterized as:

$$x_{ij} = \beta_{age} + \mu$$

The phenotypic data, which were constrained to unity, were parameterized as:

$$\sigma^2 = \sigma_A^2 + \sigma_D^2 + \sigma_E^2 \text{ or, } \sigma^2 = \sigma_A^2 + \sigma_C^2 + \sigma_E^2$$

where, $\sigma^2_A$ represents additive genetic effect (A); $\sigma^2_D$ represents non-additive genetic effects (D); $\sigma^2_C$ represents shared environmental effects(C) and $\sigma^2_E$ represents non-shared or unique environmental effects (E). The covariance terms were parameterized as:

$$Cov_{MZs} = \sigma_A^2 + \sigma_D^2 \text{ or } \sigma_A^2 + \sigma_C^2$$

$$Cov_{DZs} = .5\sigma_A^2 + .25\sigma_D^2 \text{ or } .5\sigma_A^2 + \sigma_C^2$$

The significance of variance components was tested by comparing the fit (minus twice the log-likelihood) of the full model which included the effect to that of a nested model in which the effect had been dropped from the model. The difference in log-likelihoods follows an asymptotic chi-square distribution with the degrees of freedom equal to the difference in estimated parameters between the two models. The results of these analyses are summarized in **Supplementary Table 5**.

#### Look-up in GWAS browsers

In order to assess the potential genetic overlap between miscarriage phenotypes and other traits, we conducted a lookup in the publicly available GWAS catalogue (https://www.ebi.ac.uk/gwas/), and GWAS browsers Oxford BIG browser (http://big.stats.ox.ac.uk/) and GWASAtlas[26] (http://atlas.ctglab.nl/) for the miscarriage-associated variants identified in any of our three GWAS meta-analyses. One of our variants associated with recurrent miscarriage (chr 9: rs7859844 showed nominal association with 'stomach or abdominal pain' ($P$=4.9×10$^{-6}$) and venous thromboembolic disease ($P$=5.8×10$^{-6}$)[26], 'CD314-CD158a+ NK cell proportion' ($P$=3.3×10$^{-6}$) and CD32+ mDC (dendritic cell) subset proportion ($P$=8.7×10$^{-6}$)[27].

#### Genetic correlation analyses

The LD Score regression method[23] implemented in LD-Hub (http://ldsc.broadinstitute.org)[28] was used for testing genetic correlations between sporadic miscarriage and 72 traits (spanning reproductive, anthropometric, psychiatric, aging, haematological, cardiometabolic, autoimmune, hormone, cancer and smoking behaviour traits), using the sporadic miscarriage European-ancestry only GWAS meta-analysis summary statistics and data available within the LD Hub resource. Bonferroni correction ($0.05/72=6.9\times10^{-4}$) was used to account for multiple testing. Results of the analysis are presented in **Supplementary Table 6**.

**Associated phenotypes analysis**

The UKBB has extensive phenotype data for 500K individuals, of which 273,465 are women. After applying the same QC criteria that were applied to the subset of individuals used in the genetic association analysis, extensive phenotype data was available for 220,804 women. Among these there were 39,411 sporadic miscarriage cases and 133,545 controls, and 458 recurrent miscarriage cases and 133,675 controls (defined using the criteria applied to define cases and controls for the association analysis). We explored differences in the prevalence of diseases between sporadic miscarriage cases and controls and between recurrent miscarriage cases and controls. Diseases were identified from the UKBB linked Hospital Episode Statistics which provide ICD10 diagnosis codes (UKBB data fields 41202 and 41204). First, for each hospital diagnosis observed among the cases (defined by an ICD10 code; n=6,840 for sporadic and n=1,323 for recurrent miscarriage, respectively; excluding those used to define the cases), we tested the difference in the proportion of cases with the diagnosis with that among the controls (using a 2-sample test for difference in proportions when the number of "successes" and "failures" are greater or equal to five for both populations; otherwise, Fisher's exact test was applied instead). A false discovery rate (FDR) multiple testing correction at level 10% was applied to the *P*-values. For graphical representation (**Supplementary Figure 5)**, diagnosis codes were grouped and coloured by ICD10 chapters as follows: blood ("Diseases of the blood and blood-forming organs and certain disorders involving the immune mechanism", D50-D89), circulatory ("Diseases of the circulatory system", I00-I99), congenital ("Congenital malformations, deformations and chromosomal abnormalities", Q00-Q99), digestive ("Diseases of the digestive system", K00-K93), ears ("Diseases of the ear and mastoid process", H60-H95), endocrine ("Endocrine, nutritional and metabolic diseases", E00-E90), external ("External causes of morbidity and mortality", V01-Y98), eyes ("Diseases of the eye and adnexa", H00-H59), genitourinary ("Diseases of the genitourinary system", N00-N99), infection ("Certain infectious and parasitic diseases", A00-B99), injury ("Injury, poisoning and certain other consequences of external causes", S00-T98), musculoskeletal ("Diseases of the musculoskeletal system and connective tissue", M00-M99), neoplasms ("Neoplasms", C00-D48), nervous ("Diseases of the nervous system", G00-G99), other ("Factors influencing health status and contact with health services", U04-Z99), perinatal ("Certain conditions originating in the perinatal period", P00-P96), pregnancy ("Pregnancy, childbirth and the puerperium", O00-O99), psychiatric ("Mental and behavioural disorders", F00-F99), respiratory ("Diseases of the respiratory system", J00-J99), skin ("Diseases of the skin and subcutaneous tissue", L00-L99), symptoms ("Symptoms, signs and abnormal clinical and laboratory findings, not elsewhere classified", R00-R99). To rule out the confounding effect of woman's age and parity, we then conducted multivariate logistic regression adjusted for age and number of children for any diseases for which there was statistical evidence of a difference between cases and controls, and applied FDR 5% correction to the *P*-values (**Supplementary Tables 7 and 8**).

**A**

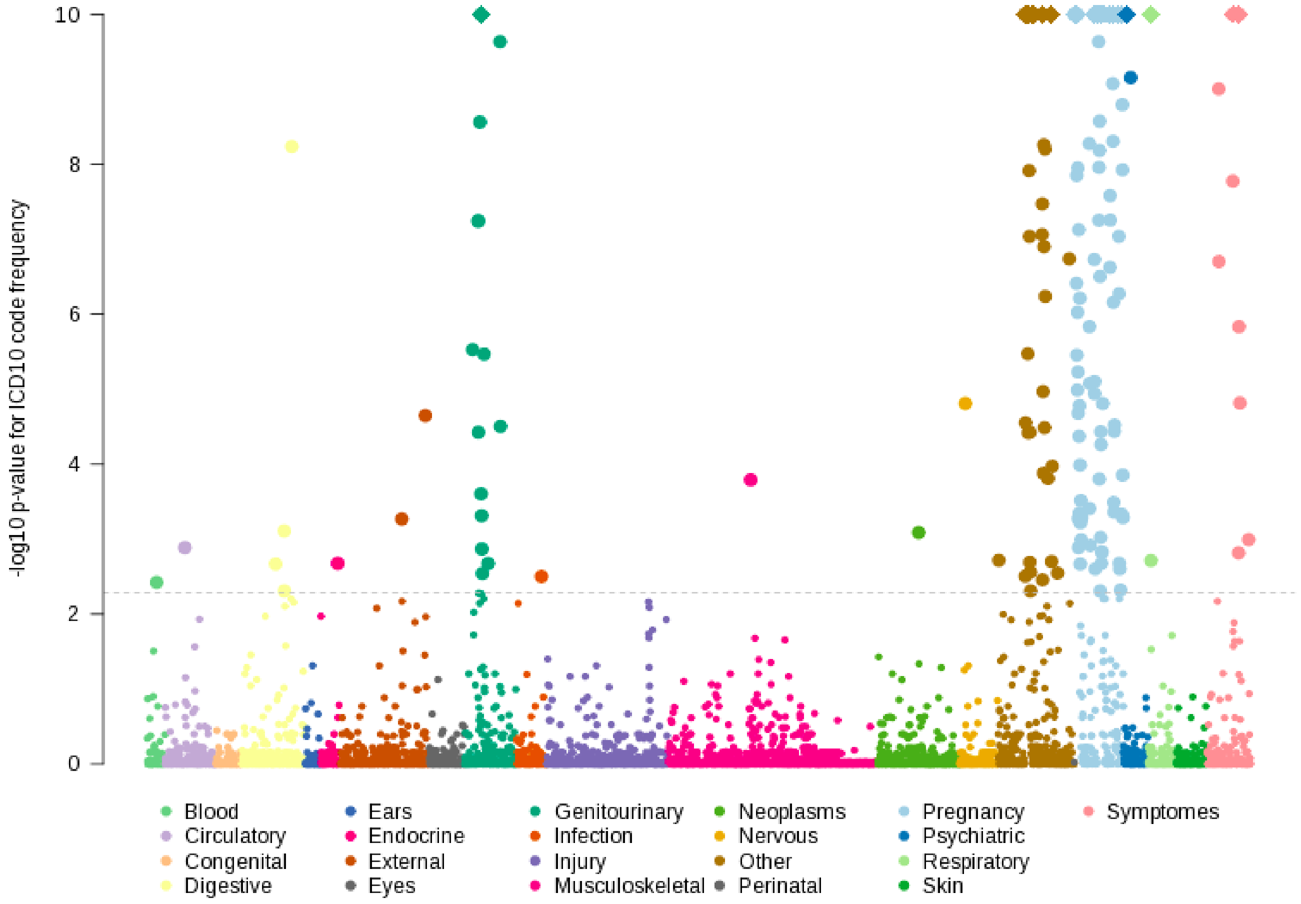

**B**

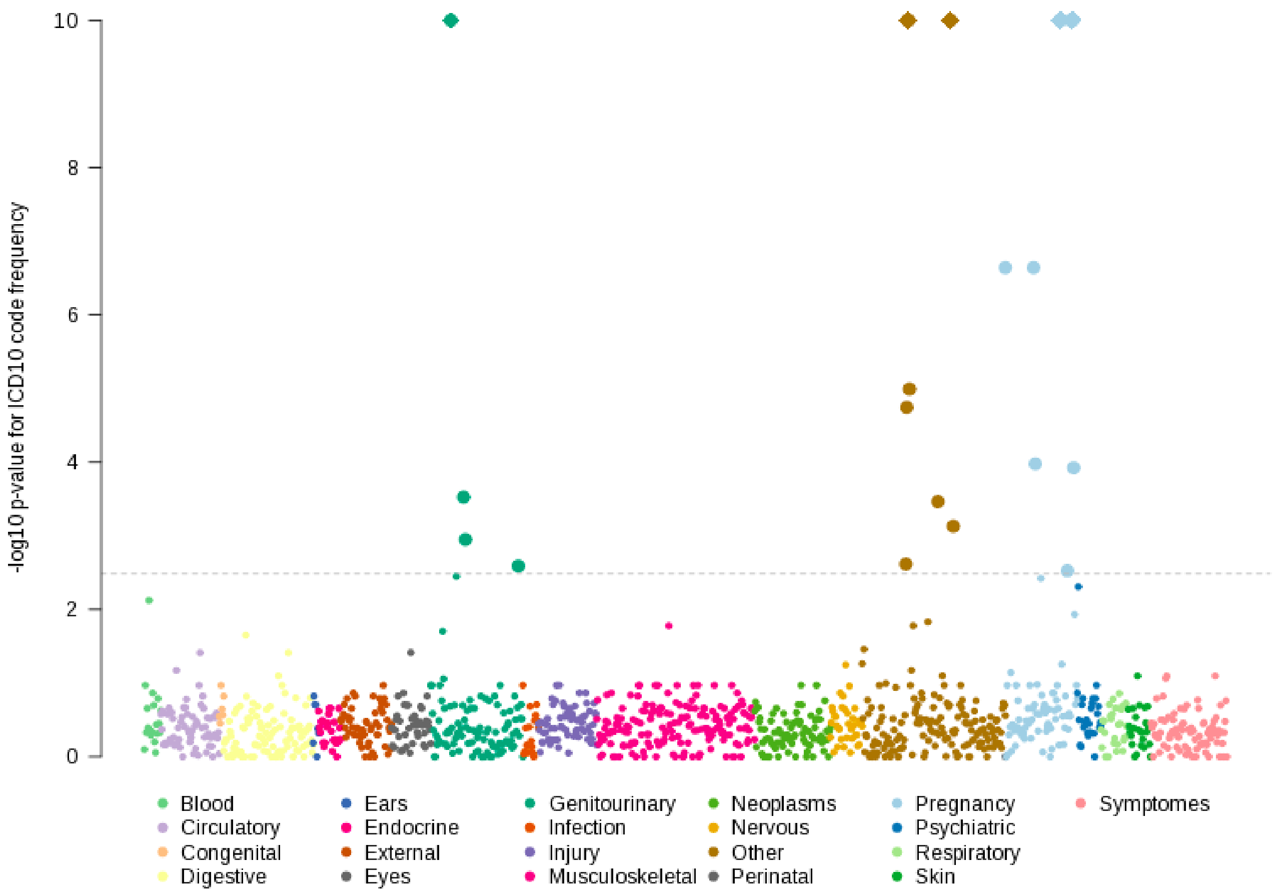

-log10 p-value for ICD10 code frequency
10
8
6
4
2
0
Blood
Circulatory
Congenital
Digestive
Ears
Endocrine
External
Eyes
Genitourinary
Infection
Injury
Musculoskeletal
Neoplasms
Nervous
Other
Perinatal
Pregnancy
Psychiatric
Respiratory
Skin
Symptomes

**Supplementary Figure 5. ICD codes associated with sporadic (A) and recurrent (B) miscarriage (unadjusted analysis).** Each point in the plot represents one ICD10 code observed among the cases. Y-axis is the –log10 *P*-value of the test of difference between the diagnosis frequency among cases and controls. Diagnoses were grouped and coloured by the ICD10 chapter. Y-axis is truncated at 10, with diagnosis for which its –log10 *P*-value exceeds this value represented by a diamond. Horizontal dash line represents the significance threshold after applying the 10% FDR multiple testing correction ($5.23\times10^{-3}$ for sporadic and $3.28\times10^{-3}$ for recurrent miscarriage, respectively).

**Mendelian randomization (MR) analyses**

We conducted a (hypothesis generating) MR phenome-wide (PheWAS) analysis of recurrent miscarriage (using a per allele genetic risk score from the four GWAS significant SNPs; rs7859844, rs143445068, rs138993181, rs183453668) in relation to 17,037 outcomes using the PHESANT[29] package in UKBB (n=168,763) (**Supplementary Figure 6**). The analysis was adjusted for year of birth and the top 10 PCs. Overall, the MR-PheWAS did not show any evidence of causal effects (**Supplementary Figure 7**). Only 3 outcomes reached Bonferroni corrected levels of statistical significance ($P<2.93\times10^{-6}$), including one outcome related to alcoholism and one related to post-traumatic stress disorder. However, both of these were single items from instruments that included 11 items (alcohol use questionnaire) and 21 items (post-traumatic/traumatic event questionnaire), respectively, with none of the other items reaching suggestive thresholds of statistical significance. The third outcome to show association below this p-value threshold was a job coding (scenery designer or costume designer) that is one of a which lies in 42-item employment history category (MR analyses did not suggest effects on any other jobs in this list).

*Hypothesis generating and need for replication*

The nature of PheWAS are that they are hypothesis generating. Whilst the application of a stringent p-value threshold helps to minimise false positive results, any suggestive associations need replication in independent datasets and appropriate sensitivity analyses to explore whether the associations are likely to be causal or reflect horizontal pleiotropy. This approach is not able to conclusively determine the nature of any causal effect, thus outcomes that are statistically significantly related to the exposure of interest (here recurrent miscarriage) might be confounded. In this specific example, the MR-PheWAS was undertaken in a study that is not independent of the GWAS sample that identified the recurrent miscarriage hits (it is the largest study contributing to that GWAS) and there may be over-fitting of the data. Furthermore, with only four rare SNPs the use of MR-Egger to explore horizontal pleiotropy may be unreliable. Lastly, the proportion of variation in recurrent miscarriage explained by the genetic risk score (between 0.02% to 0.6%) is low (McFadden's adjusted R2=0.0006, Efron's R2=0.0002, Pseudo R2=0.0062), suggesting that we might have weak instrument bias. Given these considerations and the fact that the three associations that reached significance below our predefined Bonferroni threshold did not seem plausible causes or consequences of recurrent miscarriage we did not explore any of those further. We did look at other outcomes in the top 10 lowest *P*-values (i.e. the group that deviate from the QQ plot; **Supplementary Figure 7**) to see if any were more plausible. **Supplementary Table 9** lists the top 10 (lowest *P*-values) outcomes with per recurrent miscarriage allele association (SD) and *P*-value. With one exception none of these are outcomes that have been shown in previously published, or our, multivariable regression analyses and do not seem biologically plausible. The one exception is a suggestive causal effect of recurrent miscarriage on endometriosis of the uterus ($P$=5.9×10$^{-5}$).

**Two-sample Mendelian randomization (MR) of endometriosis on recurrent miscarriage**

We used a two-sample MR approach[30] to explore the possible causal effect of endometriosis on recurrent miscarriage. We obtained summary association results for genetic instruments for endometriosis from a recent meta-analysis of 11 genome-wide association case-control data sets[31]. Summary associations between each instrument and recurrent miscarriage were estimated using Firth regression in the three European ancestry studies of our GWAS and meta-analysed using METAL[19]. From the 19 SNPs associated with endometriosis, 18 were present in at least two of our three studies. After harmonization of the summary data sets, we used inverse variance weighting (IVW)[32] and MR-Egger[33] to obtain a pooled estimate of the association between the SNPs for endometriosis and recurrent miscariage. There was no evidence of a causal association between endometriosis and recurrent miscarriage (IVW OR 1.18, 95% CI 0.84, 1.65 and MR-Egger OR 0.96, 95% CI 0.23, 4.02) (**Supplementary Figure 8**).

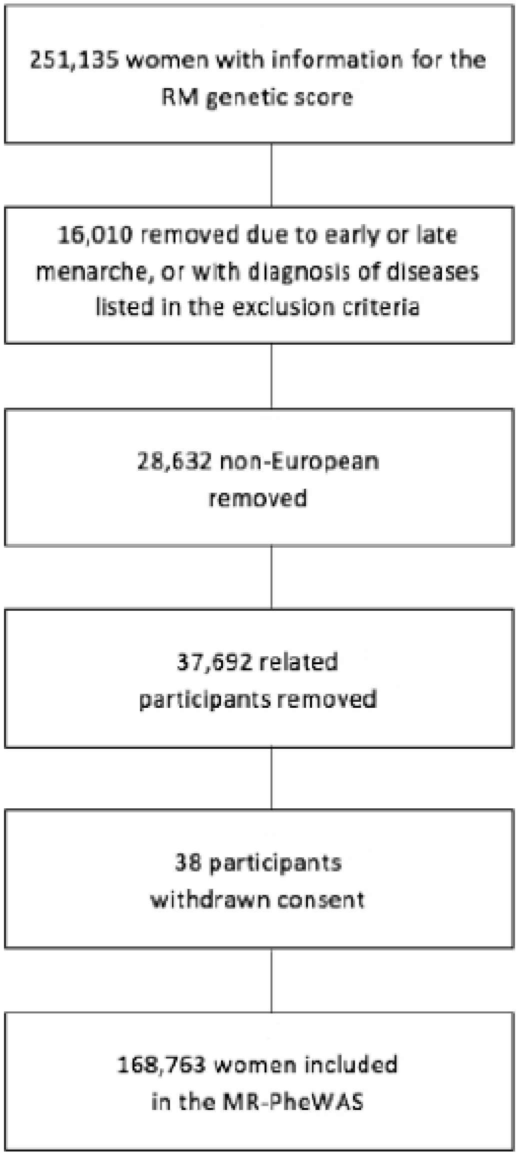


**Supplementary Figure 6. Participant flow diagram for the MR-PheWAS**

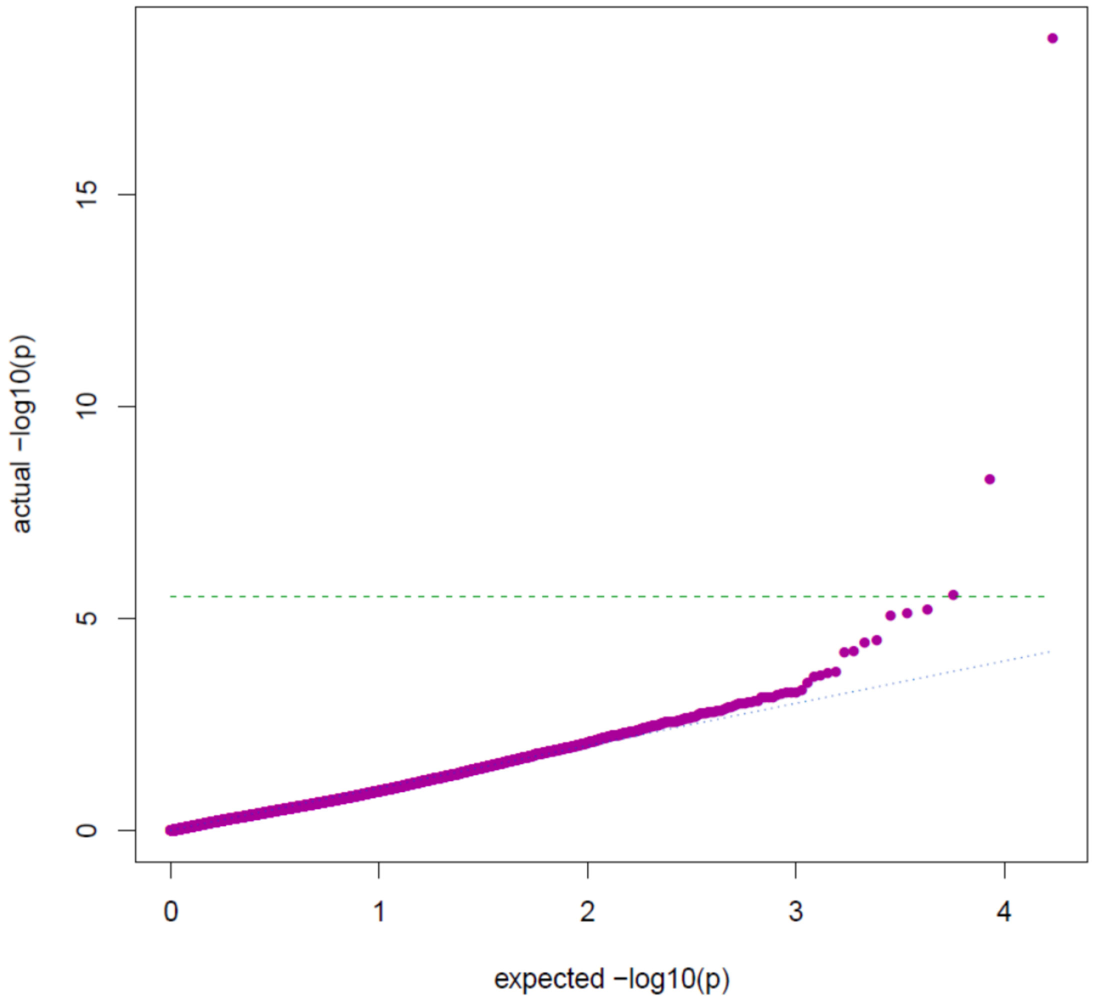


**Supplementary Figure 7.** QQ plot for the 17,028 MR-PheWAS results for recurrent miscarriage genetic risk score.

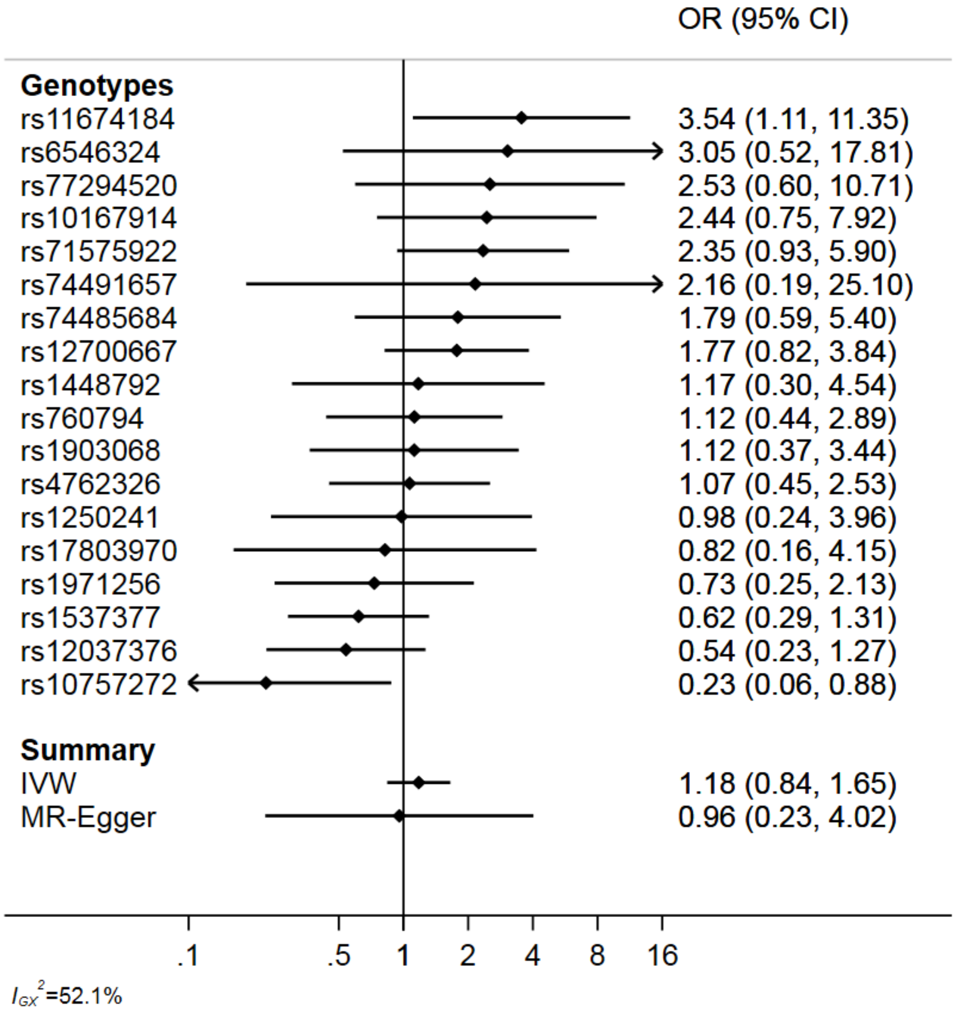


**Supplementary Figure 8.** Forest plot showing risk allele effects for recurrent miscarriage and pooled effect according to inverse variance weighting and MR-Egger methods.

**Functional annotation**

The FUMA platform designed for prioritization, annotation and interpretation of GWAS results[21] was used for functional annotation of association signals from the GWAS meta-analyses. As the first step, independent significant SNPs in the GWAS meta-analysis summary statistics were identified based on their P-values ($P < 5\times10^{-8}$) and independence from each other (r2<0.6 in the 1000G phase 3 reference) within a 1Mb window. Thereafter, lead SNPs were identified from independent significant SNPs, which are independent of each other (r2<0.1). SNPs that were in LD with the identified independent SNPs (r2≥0.6) within a 1Mb window, have a MAF of ≥1% and GWAS meta-analysis *P*-value of >0.05 were selected as candidate SNPs and taken forward for further annotation.

FUMA annotates candidate SNPs in genomic risk loci based on functional consequences on genes using the Annotate Variation (ANNOVAR)[34], CADD (a continuous score showing how deleterious the SNP is to protein structure/function; scores >12.37 indicate potential pathogenicity)[35] and RegulomeDB[36] scores (ranging from 1 to 7, where lower score indicates greater evidence for having regulatory function), 15 chromatin states from the Roadmap Epigenomics Project[37,38], eQTL data (GTEx v6 and v7)[39], blood eQTL browser[40], BIOS QTL browser[41], BRAINEAC[42], MuTHER[43], xQTLServer[44], the CommonMind Consortium[45], and 3D chromatin interactions from HI-C experiments of 21 tissues/cell types[46]. FUMA maps genes to candidate SNPs using positional mapping, which is based on ANNOVAR annotations and maximum distance between SNPs (default 10 kb) and genes, eQTL mapping and chromatin interaction mapping. Chromatin interaction mapping was performed with significant chromatin interactions (defined as FDR<$1\times10^{-6}$). The two ends of significant chromatin interactions were defined as follows: region 1, a region overlapping with one of the candidate SNPs; and region 2, another end of the significant interaction, used to map to genes based on overlap with a promoter region (250 bp upstream and 50 bp downstream of the transcription start site).

To narrow down potential candidate genes, we used Hi-C chromatin interaction datasets to visualize topologically associated domains (TADs) in the region and Capture Hi-C data for various tissues to further explore interactions within the TAD domain. Data was visualised using the 3D Genome Browser[47] (http://3dgenome.org). TADs are relatively conserved across different tissue types and define the boundaries for potential genomic interactions[48].

For sporadic miscarriage, we used the summary statistics of our EUR-ancestry meta-analysis. A total of five candidate SNPs were identified ($r^2$>=0.6 with rs146350366) in the associated locus on chr13, all of them intergenic (**Supplementary Table 10**). Of these, rs188519103, located 6.9kb 5' of *SNORD36* had the lowest RegulomeDB score (4 - evidence of transcription factor binding and DNase peak). Potential candidate genes were mapped using eQTL and chromatin interaction data (**Supplementary Table 11**).

In the recurrent miscarriage analysis, we had 3 associated loci with consistent effect direction in all three cohorts. For the signal on chromosome 9, 53 candidate SNPs were identified by FUMA (**Supplementary Table 12**). Of these, rs12004880 had a RegulomeDB score of 3a ("TF binding + any motif + DNase peak"), while four SNPs had a CADD score of>12.37, indicating potential pathogenicity[35]. A total of 50 candidate genes were proposed (**Supplementary Table 13**), among them protein-coding *TLE1, TLE4, PSAT1, IDNK, GNAQ, RASEF, SPATA31D1* and *FRMD3*. On chromosome 11, rs143445068 (RegulomeDB score 3a) and rs140847838 were highlighted as potential candidate SNPs in the associated region located in the intron of *NAV2*. Chromatin interaction mapping proposed another 17 candidate genes, including *DBX1, HTATIP2, E2F8, ZDHHC13, MRGPRX2*. Hi-C map in ovaries from the 3D Genome Browser[47] is shown on **Supplementary Figure 9**. Finally, for association signal on chromosome 21, no other candidate SNPs in addition to the lead signal rs183453668 were identified, and a total of 10 candidate genes were suggested by chromatin interaction data. Hi-C map in ovaries and Capture Hi-C data visualization in endothelial progenitors are shown on **Supplementary Figure 10**.

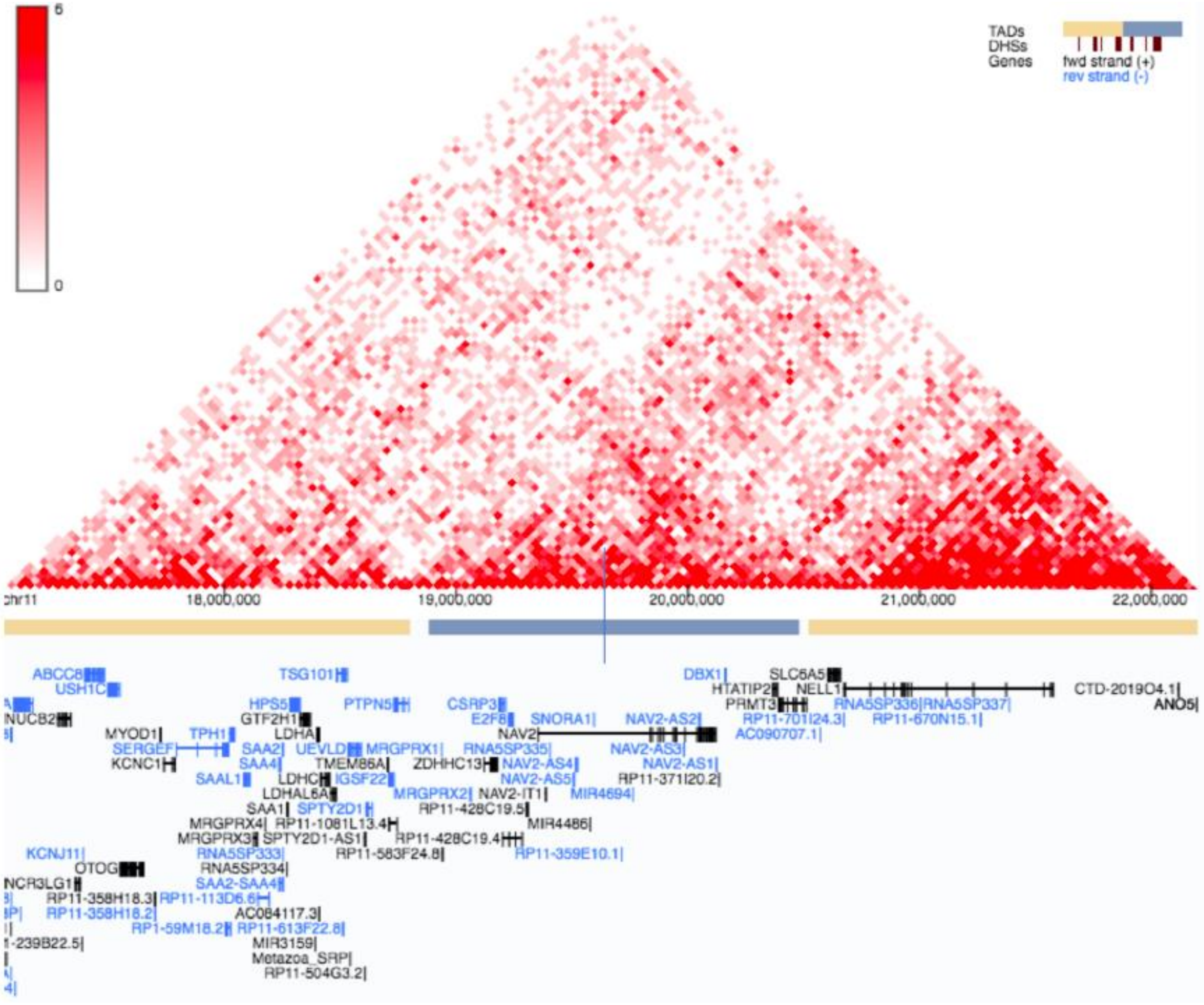


**Supplementary Figure 9. Hi-C map in ovaries for the recurrent miscarriage association signal on chromosome 11.** The blue vertical line represents the location of the signal from GWAS meta-analysis. The 3D Genome Browser[47] was used for data visualization.

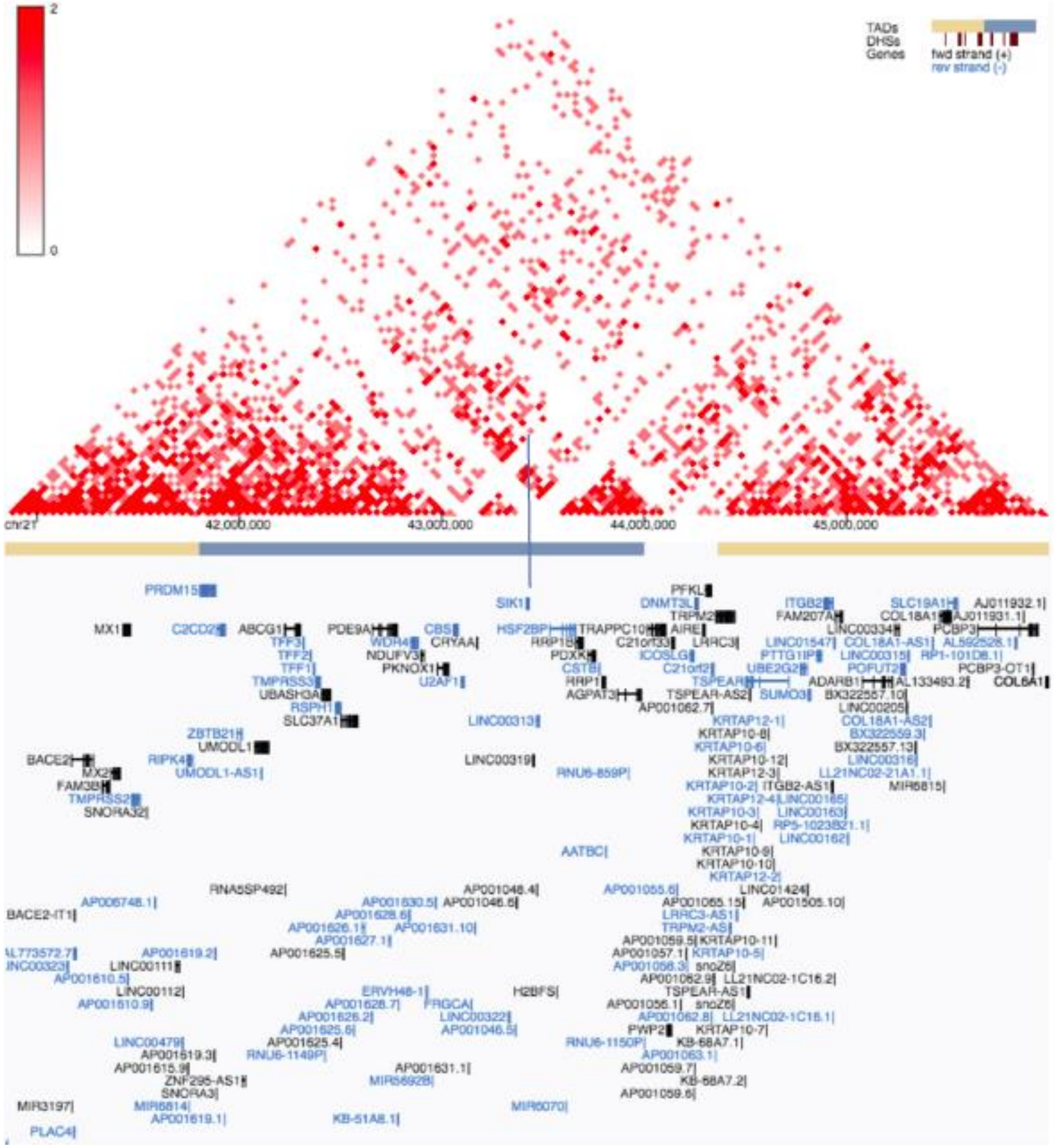
2
0
TADs
DHSs
Genes
fwd strand (+)
rev strand (-)
chr21
42,000,000
43,000,000
44,000,000
45,000,000

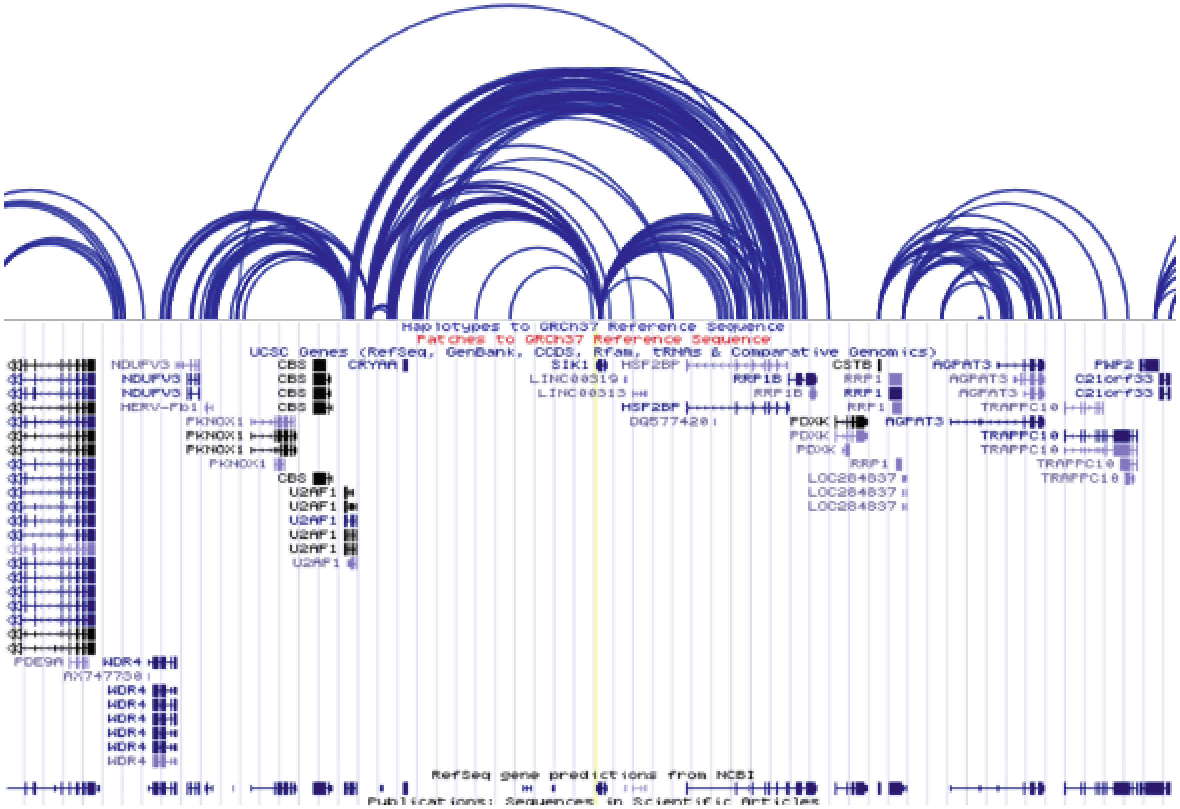


**Supplementary Figure 10. Hi-C map in ovaries and Capture Hi-C data visualization in endothelial progenitors for recurrent miscarriage association signal on chr21.** The blue/yellow vertical line represents the location of the signal from GWAS meta-analysis. The 3D Genome Browser[47] was used for data visualization.

**Acknowledgements**

T.L. is supported by European Commission Horizon 2020 research and innovation programme (project WIDENLIFE, grant number 692065); Estonian Ministry of Education and Research (grants IUT34-16 and PUTJD726); Enterprise Estonia (grant EU49695). C.M.L. is supported by the Li Ka Shing Foundation, WT-SSI/John Fell funds, Oxford, NIHR Oxford Biomedical Research Centre, Oxford, Widenlife and NIH (5P50HD028138-27). D.F.C. is supported by grants from the National Institutes of Health (R01HD078641, R01MH101810 and P51OD011092). T.F. is supported by the NIHR Biomedical Research Centre, Oxford.

*UKBB*

The research in this paper has been carried out using the UK Biobank resource (under applications 17805 ("Dissemination of shared genetics across phenotypes associated with reproductive health and related endophenotypes"), 11867 ("Dissection of the Genetic Susceptibility of Obesity Traits and their Comorbidities"), and 16729 ("MR-PheWAS: hypothesis prioritization among potential causal effects of body mass index on many outcomes, using Mendelian randomization"))

*EGCUT*

This study was funded by EU H2020 grant 692145, Estonian Research Council Grant IUT20-60, IUT24-6, and European Union through the European Regional Development Fund Project No. 2014-2020.4.01.15-0012 GENTRANSMED and 2014-2020.4.01.16-0125. Data analyses were carried out in part in the High-Performance Computing Center of University of Tartu.

*iPSYCH*

iPSYCH2012 has been conducted using the Danish National Biobank resource, supported by the Novo Nordisk Foundation.

*QIMR*

We thank the twins and their families for their participation in the QIMR study. Funding was provided by the Australian National Health and Medical Research Council (241944, 339462, 389927, 389875, 389891, 389892, 389938, 442915, 442981, 496739, 552485, 552498, 1084325), the Australian Research Council (A7960034, A79906588, A79801419, DP0770096, DP0212016, DP0343921), the FP-5

GenomEUtwin Project (QLG2-CT-2002-01254), and the U.S. National Institutes of Health (NIH grants AA07535, AA10248, AA13320, AA13321, AA13326, AA14041, MH66206). S.E.M. is supported by a National Health and Medical Research Council (NHMRC) Senior Research Fellowship (1103623).

*QIMREndo*

We acknowledge with appreciation all women who participated in the QIMR endometriosis study. This study was supported by grants from the Australian National Health and Medical Research Council (241944, 339462, 389927,389875, 389891, 389892, 389938, 443036, 442915, 442981, 496610, 496739, 552485, 552498), the Cooperative Research Centre for Discovery of Genes for Common Human Diseases (CRC), Cerylid Biosciences (Melbourne), and donations from Neville and Shirley Hawkins. D.R.N and G.W.M are supported by the NHMRC Fellowship Scheme.

*ALSPAC*

We are extremely grateful to all the families who took part in the ALSPAC study, the midwives for their help in recruiting them, and the whole ALSPAC team, which includes interviewers, computer and laboratory technicians, clerical workers, research scientists, volunteers, managers, receptionists and nurses. FUNDING: This study was supported by the US National Institute of Health (R01 DK10324), which also supports A.L.G.S' salary, the European Research Council under the European Union's Seventh Framework Programme (FP7/2007-2013) / ERC grant agreement no 669545 and the NIHR Biomedical Centre at the University Hospitals Bristol NHS Foundation Trust and the University of Bristol. Core funding for ALSPAC is provided by the UK Medical Research Council and Wellcome (Grant ref: 102215/2/13/2) and the University of Bristol. Genotyping of the ALSPAC maternal samples was funded by the Wellcome Trust (WT088806). A comprehensive list of grants funding is available on the ALSPAC website (http://www.bristol.ac.uk/alspac/external/documents/grant-acknowledgements.pdf). A.L.G.S and D.A.L work in a Unit that receives support from the University of Bristol and UK Medical Research Council (MC_UU_00011/6). None of the funders influenced the analyses or interpretation of results. The views expressed in this paper are those of the authors and not necessarily any funding body. D.A.L and A.L.G.S will serve as guarantors for the results from ALSPAC.

*WHI*

The WHI program is funded by the National Heart, Lung, and Blood Institute, National Institutes of Health, U.S. Department of Health and Human Services through contracts HHSN268201600018C, HHSN268201600001C, HHSN268201600002C, HHSN268201600003C, and HHSN268201600004C. This manuscript was not prepared in collaboration with investigators of the WHI, has not been reviewed and/or approved by the Women's Health Initiative (WHI), and does not necessarily reflect the opinions of the WHI investigators or the NHLBI. Funding support for WHI GARNET was provided through the NHGRI Genomics and Randomized Trials Network (GARNET) (Grant Number U01 HG005152). Assistance with phenotype harmonization and genotype cleaning, as well as with general study coordination, was provided by the GARNET Coordinating Center (U01 HG005157). Assistance with data cleaning was provided by the National Center for Biotechnology Information. Funding support for genotyping, which was performed at the Broad Institute of MIT and Harvard, was provided by the NIH Genes, Environment and Health Initiative [GEI] (U01 HG004424). Funding for WHI SHARe genotyping was provided by NHLBI Contract N02-HL-64278. The WHI datasets used for the analyses described in this manuscript were obtained from dbGaP at http://www.ncbi.nlm.nih.gov/sites/entrez?db=gap through dbGaP accessions phs000200.v11.p3.c1 and phs000200.v11.p3.c2.

*Partners HealthCare Biobank*

Partners HealthCare Biobank is supported by NHGRI grant U01HG008685.

*Lifelines*

The Lifelines Cohort Study, and generation and management of GWAS genotype data for the Lifelines Cohort Study is supported by the Netherlands Organization of Scientific Research NWO (grant 175.010.2007.006), the Economic Structure Enhancing Fund (FES) of the Dutch government, the Ministry of Economic Affairs, the Ministry of Education, Culture and Science, the Ministry for Health, Welfare and Sports, the Northern Netherlands Collaboration of Provinces (SNN), the Province of Groningen, University Medical Center Groningen, the University of Groningen, Dutch Kidney Foundation and Dutch Diabetes Research Foundation.

MoBa

This work was supported by grants from the European Research Council (AdG #293574), the Bergen Research Foundation ("Utilizing the Mother and Child Cohort and the Medical Birth Registry for Better Health"), Stiftelsen Kristian Gerhard Jebsen (Translational Medical Center), the University of Bergen, the Research Council of Norway (FRIPRO grant #240413), the Western Norway Regional Health Authority (Strategic Fund "Personalized Medicine for Children and Adults"), and the Norwegian Diabetes Foundation; and Helse Vest's Open Research Grant. This work was partly supported by the Research Council of Norway through its Centres of Excellence funding scheme (#262700), Better Health by Harvesting Biobanks (#229624) and The Swedish Research Council, Stockholm, Sweden (2015-02559), The Research Council of Norway, Oslo, Norway (FRIMEDBIO ES547711, March of Dimes (#21-FY16-121). The Norwegian Mother and Child Cohort Study is supported by the Norwegian Ministry of Health and Care Services and the Ministry of Education and Research, NIH/NIEHS (contract no N01-ES-75558), NIH/NINDS (grant no.1 UO1 NS 047537-01 and grant no.2 UO1 NS 047537-06A1). We are grateful to all the families in Norway who are taking part in this ongoing cohort study.

Kadoorie Biobank

The CKB baseline survey and the first re-survey were supported by the Kadoorie Charitable Foundation in Hong Kong. Long-term follow-up was supported by the UK Wellcome Trust (202922/Z/16/Z, 104085/Z/14/Z, 088158/Z/09/Z), the National Natural Science Foundation of China (81390540, 81390541, 81390544), and the National Key Research and Development Program of China (2016YFC0900500, 2016YFC0900501, 2016YFC0900504, 2016YFC1303904). DNA extraction and genotyping were supported by GlaxoSmithKline and the UK Medical Research Council (MC-PC-13049, MC-PC-14135). The project was supported by British Heart Foundation, UK Medical Research Council and Cancer Research provide core funding to the Clinical Trial Service Unit and Epidemiological Studies Unit at Oxford University.

1. Sudlow, C. *et al.* UK Biobank: An Open Access Resource for Identifying the Causes of a Wide Range of Complex Diseases of Middle and Old Age. *PLoS Med.* **12,** e1001779 (2015).
2. Leitsalu, L. *et al.* Cohort Profile: Estonian Biobank of the Estonian Genome Center, University of Tartu. *Int. J. Epidemiol.* **44,** 1137–1147 (2015).
3. Fraser, A. *et al.* Cohort Profile: the Avon Longitudinal Study of Parents and Children: ALSPAC mothers cohort. *Int. J. Epidemiol.* **42,** 97–110 (2013).
4. Medland, S. E. *et al.* Males Do Not Reduce the Fitness of Their Female Co-Twins in Contemporary Samples. *Twin Res. Hum. Genet.* **11,** 481–487 (2008).
5. Painter, J. N. *et al.* Genome-wide association study identifies a locus at 7p15.2 associated with endometriosis. *Nat. Genet.* **43,** 51–54

(2011).
6. Treloar, S. A. *et al.* Genomewide linkage study in 1,176 affected sister pair families identifies a significant susceptibility locus for endometriosis on chromosome 10q26. *Am. J. Hum. Genet.* **77,** 365–76 (2005).
7. Stolk, R. P. *et al.* Universal risk factors for multifactorial diseases: LifeLines: a three-generation population-based study. *Eur. J. Epidemiol.* **23,** 67–74 (2008).
8. Scholtens, S. *et al.* Cohort Profile: LifeLines, a three-generation cohort study and biobank. *Int. J. Epidemiol.* **44,** 1172–80 (2015).
9. Rønningen, K. S. *et al.* The biobank of the Norwegian mother and child cohort Study: A resource for the next 100 years. *Eur. J. Epidemiol.* **21,** 619–625 (2006).
10. Magnus, P. *et al.* Cohort Profile Update: The Norwegian Mother and Child Cohort Study (MoBa). *Int. J. Epidemiol.* **45,** 382–388 (2016).
11. Magnus, P. *et al.* Cohort profile: The Norwegian Mother and Child Cohort Study (MoBa). *Int. J. Epidemiol.* **35,** 1146–1150 (2006).
12. Liu, S. *et al.* Genomic Analyses from Non-invasive Prenatal Testing Reveal Genetic Associations, Patterns of Viral Infections, and Chinese Population History. *Cell* **175,** 347–359.e14 (2018).
13. Chen, Z. *et al.* China Kadoorie Biobank of 0.5 million people: survey methods, baseline characteristics and long-term follow-up. *Int. J. Epidemiol.* **40,** 1652–66 (2011).
14. Winkler, T. W. *et al.* Quality control and conduct of genome-wide association meta-analyses. *Nat. Protoc.* **9,** 1192–1212 (2014).
15. McCarthy, S. *et al.* A reference panel of 64,976 haplotypes for genotype imputation. *Nat. Genet.* **48,** 1279–1283 (2016).
16. 1000 Genomes Project Consortium *et al.* A global reference for human genetic variation. *Nature* **526,** 68–74 (2015).
17. Mägi, R. *et al.* Trans-ethnic meta-regression of genome-wide association studies accounting for ancestry increases power for discovery and improves fine-mapping resolution. *Hum. Mol. Genet.* **26,** 3639–3650 (2017).
18. Loh, P.-R., Kichaev, G., Gazal, S., Schoech, A. P. & Price, A. L. Mixed-model association for biobank-scale datasets. *Nat. Genet.* **50,** 906–908 (2018).
19. Willer, C. J., Li, Y. & Abecasis, G. R. METAL: fast and efficient meta-analysis of genomewide association scans. *Bioinformatics* **26,** 2190–2191 (2010).
20. de Leeuw, C. A., Mooij, J. M., Heskes, T. & Posthuma, D. MAGMA: generalized gene-set analysis of GWAS data. *PLoS Comput. Biol.* **11,** e1004219 (2015).

21. Watanabe, K., Taskesen, E., van Bochoven, A. & Posthuma, D. Functional mapping and annotation of genetic associations with FUMA. *Nat. Commun.* **8,** 1826 (2017).
22. Pereza, N., Ostojić, S., Kapović, M. & Peterlin, B. Systematic review and meta-analysis of genetic association studies in idiopathic recurrent spontaneous abortion. *Fertil. Steril.* **107,** 150–159.e2 (2017).
23. Bulik-Sullivan, B. *et al.* An atlas of genetic correlations across human diseases and traits. *Nat. Genet.* **47,** 1236–41 (2015).
24. Neale, M. C. & Cardon, L. R. *Methodology for Genetic Studies of Twins and Families*. (Springer Netherlands, 1992). doi:10.1007/978-94-015-8018-2
25. Neale, M. C., Boker, S. M., Xie, G. & Maes, H. H. *Mx: Statistical Modeling (6th Edition ed.)*. (2006). at <http://www.vcu.edu/mx/>
26. Watanabe, K. *et al.* A global view of pleiotropy and genetic architecture in complex traits. *bioRxiv* 500090 (2018). doi:10.1101/500090
27. Roederer, M. *et al.* The genetic architecture of the human immune system: a bioresource for autoimmunity and disease pathogenesis. *Cell* **161,** 387–403 (2015).
28. Zheng, J. *et al.* LD Hub: a centralized database and web interface to perform LD score regression that maximizes the potential of summary level GWAS data for SNP heritability and genetic correlation analysis. *Bioinformatics* btw613 (2016). doi:10.1093/bioinformatics/btw613
29. Millard, L. A. C., Davies, N. M., Gaunt, T. R., Davey Smith, G. & Tilling, K. Software Application Profile: PHESANT: a tool for performing automated phenome scans in UK Biobank. *Int. J. Epidemiol.* **47,** 29–35 (2017).
30. Pierce, B. L. & Burgess, S. Efficient design for Mendelian randomization studies: subsample and 2-sample instrumental variable estimators. *Am. J. Epidemiol.* **178,** 1177–84 (2013).
31. Sapkota, Y. *et al.* Meta-analysis identifies five novel loci associated with endometriosis highlighting key genes involved in hormone metabolism. *Nat. Commun.* **8,** 15539 (2017).
32. Burgess, S., Butterworth, A. & Thompson, S. G. Mendelian randomization analysis with multiple genetic variants using summarized data. *Genet. Epidemiol.* **37,** 658–65 (2013).
33. Bowden, J., Davey Smith, G. & Burgess, S. Mendelian randomization with invalid instruments: effect estimation and bias detection through Egger regression. *Int. J. Epidemiol.* **44,** 512–25 (2015).
34. Wang, K., Li, M. & Hakonarson, H. ANNOVAR: functional annotation of genetic variants from high-throughput sequencing data.

*Nucleic Acids Res.* **38,** e164 (2010).
35. Kircher, M. *et al.* A general framework for estimating the relative pathogenicity of human genetic variants. *Nat. Genet.* **46,** 310–5 (2014).
36. Boyle, A. P. *et al.* Annotation of functional variation in personal genomes using RegulomeDB. *Genome Res.* **22,** 1790–7 (2012).
37. Dunham, I. *et al.* An integrated encyclopedia of DNA elements in the human genome. *Nature* **489,** 57–74 (2012).
38. Roadmap Epigenomics Consortium, A. *et al.* Integrative analysis of 111 reference human epigenomes. *Nature* **518,** 317–30 (2015).
39. Lonsdale, J. *et al.* The Genotype-Tissue Expression (GTEx) project. *Nat. Genet.* **45,** 580–585 (2013).
40. Westra, H.-J. *et al.* Systematic identification of trans eQTLs as putative drivers of known disease associations. *Nat. Genet.* **45,** 1238–1243 (2013).
41. Zhernakova, D. V *et al.* Identification of context-dependent expression quantitative trait loci in whole blood. *Nat. Genet.* **49,** 139–145 (2016).
42. Ramasamy, A. *et al.* Genetic variability in the regulation of gene expression in ten regions of the human brain. *Nat. Neurosci.* **17,** 1418–1428 (2014).
43. Grundberg, E. *et al.* Mapping cis- and trans-regulatory effects across multiple tissues in twins. *Nat. Genet.* **44,** 1084–1089 (2012).
44. Ng, B. *et al.* An xQTL map integrates the genetic architecture of the human brain's transcriptome and epigenome. *Nat. Neurosci.* **20,** 1418–1426 (2017).
45. Fromer, M. *et al.* Gene expression elucidates functional impact of polygenic risk for schizophrenia. *Nat. Neurosci.* **19,** 1442–1453 (2016).
46. Schmitt, A. *et al.* A Compendium of Chromatin Contact Maps Reveals Spatially Active Regions in the Human Genome. *Cell Rep.* **17,** 2042–2059 (2016).
47. Wang, Y. *et al.* The 3D Genome Browser: a web-based browser for visualizing 3D genome organization and long-range chromatin interactions. *Genome Biol.* **19,** 151 (2018).
48. Dixon, J. R. *et al.* Chromatin architecture reorganization during stem cell differentiation. *Nature* **518,** 331–336 (2015).
